# B-cell epitope mapping of TprC and TprD variants of *Treponema pallidum* subspecies

**DOI:** 10.1101/2022.01.26.477776

**Authors:** Barbara Molini, Mark C. Fernandez, Charmie Godornes, Anastassia Vorobieva, Sheila A. Lukehart, Lorenzo Giacani

## Abstract

Several recent studies have focused on the identification, functional analysis, and structural characterization of outer membrane proteins (OMPs) of *Treponema pallidum* (*Tp*). The *Tp* species encompasses the highly related *pallidum*, *pertenue,* and *endemicum* subspecies of this pathogen, known to be the causative agents of syphilis, yaws, and bejel, respectively. These studies highlighted the importance of identifying surface-exposed OMP regions and the identification of B-cell epitopes that could be protective and used in vaccine development efforts. We previously reported that the TprC and TprD OMPs of *Tp* are predicted to contain external loops scattered throughout the entire length of the proteins, several of which show a low degree of sequence variability among strains and subspecies. In this study, these models were corroborated using AlphaFold2, a state-of-the-art protein structure modeling software. Here, we identified B-cell epitopes across the full-length TprC and TprD variants using the Geysan pepscan mapping approach with antisera from rabbits infected with syphilis, yaws, and bejel strains and from animals immunized with refolded recombinant TprC proteins from three syphilis strains. Our results show that the humoral response is primarily directed to sequences predicted to be on surface-exposed loops of TprC and TprD proteins, and that the magnitude of the humoral response to individual epitopes differs among animals infected with various syphilis strains and *Tp* subspecies. Rather than exhibiting strain-specificity, antisera showed various degrees of cross-reactivity with variant sequences from other strains. The data support the further exploration of TprC and TprD as vaccine candidates.

## INTRODUCTION

The human treponematoses (syphilis, yaws, and bejel) are caused by a group of highly related pathogens classified as subspecies of the spirochete bacterium *Treponema pallidum* (*Tp*). Classically, the *pallidum* subspecies is said to causes syphilis, while the *pertenue* and *endemicum* subspecies are regarded as the causes of yaws and bejel, respectively (1), although the modes of transmission and the clinical manifestations are similar among subspecies. These diseases are still a concern for public and global health, as they continue to result in substantial morbidity and mortality worldwide. According to the World Health Organization, the global prevalence of syphilis is ∼20 million cases, with an incidence of ∼6.3 million new cases every year (2). Although most of these infections occur in low- and middle-income countries, syphilis has resurged in industrialized nations of Asia, Europe, and North America. In the United States, the incidence of infectious syphilis has risen steadily over the last two decades (3–7) reaching approximately 39,000 cases in 2019, a 6.5-fold increase compared to the ∼6,000 cases reported in 2000. If left untreated, syphilis can progress to affect the cardiovascular and central nervous systems of patients, potentially leading to manifestations such as aortic aneurysm, stroke, hearing or visual loss, dementia, paralysis, and death (8). Additionally, vertical transmission of syphilis is estimated to account for ∼1/3 of stillbirths in sub-Saharan Africa and a high proportion of perinatal morbidity and mortality globally (9, 10). Past public health initiatives to eliminate syphilis and congenital syphilis promoted by the CDC and WHO (11, 12) have significantly aided in reducing syphilis incidence and in generating awareness of this disease, but have not achieved their intended elimination goals. Compared to syphilis, less accurate epidemiological data are available on yaws and bejel. Although it was recently estimated that ∼65,000 cases of yaws occurred annually in 13 endemic countries, this is likely an underestimate of the global burden of the disease, given that in at least 19 potentially endemic countries the incidence of yaws is unknown (13). While the ongoing elimination campaign in Asia and Africa using mass administration of azithromycin has demonstrated promising results (14), such efforts could be undermined by the spreading of macrolide resistant *Tp* subsp. *pertenue,* as recently demonstrated in Papua New Guinea (15). Foci of bejel have been reported in the last two decades, mostly in the Near East and Sahelian Africa (16–19), and bejel strains have recently reported to be transmitted sexually (20).

The chance of success of current and future control campaigns for all treponematoses would significantly increase if effective vaccines were available (21, 22). The most rational approach to vaccine development for these infections requires a clear understanding of the type of immune response that is protective and the identification of suitable candidate antigens to be tested in a pre-clinical animal model (21, 22). Furthermore, because there is very limited or no cross-immunity between subspecies of *Tp* and only sporadic cross-immunity between syphilis strains (23, 24), the identification of antigenic differences in potential vaccine candidates among subspecies and strains is of pivotal importance, as such differences could be key to devising a broadly protective vaccine (22). There is consensus that vaccine candidates are most likely to be found among these spirochetes’ surface-exposed antigens, such as (but not limited to) integral outer membrane proteins (OMPs). As in all dual-membrane bacteria, *Tp* integral OMPs will necessarily contain a membrane-embedded β-barrel domain composed of antiparallel β-strands joined together by loops that alternatively protrude toward the extracellular environment or the periplasm (25). Because *Tp* clearance from early lesions is dependent on opsonophagocytosis of *Tp* cells by activated macrophages (26, 27), the identification of surface-exposed epitopes that can be targeted by immunization to induce opsonic antibodies and promote macrophage activation is key to vaccine development. Such tasks, however, have been historically challenging due to the inability to steadily propagate the *Tp* subspecies *in vitro*, which was only recently achieved (28), and also because of the uncommon fragility and limited protein content of these spirochetes’ OM (29, 30). These limitations have been partially overcome by the ability to predict *in silico* OMP-encoding genes and the structure of their encoded proteins, enabling investigation using structural and functional experimental approaches (31, 32).

Among *Tp* putative OMPs identified to date, there are several members of the *<u>T</u>. <u>p</u>allidum* <u>r</u>epeat (Tpr) family of paralogous proteins, including TprC and TprD (encoded by the *tp0117* and *tp0131* genes in the reference Nichols strain, respectively) (33); these are reported to have OM localization and porin activity (34, 35). In this study, we examine the protein sequence variation in TprC and TprD among *T. pallidum* strains and subspecies, and predict, then confirm, the locations of B cell epitopes using antisera from infected and immunized rabbits. Variant specificity and cross-immunity are analyzed so that epitopes with broad coverage among strains and subspecies can be identified for future evaluation as vaccine antigens.

## RESULTS

### Sequence analysis of TprC and TprD variants

Although the TprC and TprD proteins are identical in the Nichols, Chicago, and Bal73-1 strains, allelic variants of TprC and TprD exist among syphilis strains and the three *Tp* subspecies (35, 36). Among the treponemal strains used in this study (Fig. 1), four alleles were found at the *tprD* locus, which include the reference *tprD* allele (found in the syphilis Nichols, Chicago, and Bal73-1 strains), and the *tprD_2_* allele (found in the syphilis strains MexicoA, Sea81-4, Bal3, and UW249) which encodes the TprD_2_ protein (35). Also the subsp. *pertenue* SamoaD strain and subsp. *endemicum* IraqB strains harbor a *tprD_2_* allele in the *tprD* locus, but their TprD_2_ amino acid sequences differ from the subsp. *pallidum* TprD_2_ sequence due to five amino acid substitutions scattered throughout the length of the protein (Fig.1) (35). TprD_2_ has four unique regions that differentiate it from the reference TprD sequence. These include a large central region of 110 amino acids and three smaller regions toward the COOH-terminal end of the protein (Fig.1) (35). As previously reported, the *tprC* locus of MexicoA, Sea81-4, and Bal3 encodes a TprC variant with a limited number of amino acid (aa) changes (15 aa for MexicoA, 9 aa for Sea81-4, and 9 aa for Bal3) compared to the reference TprC found in Nichols, Chicago, and Bal73-1 strains (Fig.1) (35). The TprC protein of the *pertenue* and *endemicum* strains studied here also shows limited amino acid changes compared to the reference TprC (31 aa for SamoaD and 26 aa for IraqB; Fig.1), albeit higher compared to the subsp. *pallidum* strain (35). We previously reported that TprC and TprD/D_2_ sequence variation does not occur randomly, but rather is localized in discrete variable regions (DVRs; Fig.1) (35). When TprC and TprD variants are compared (with the exclusion of TprD_2_), seven DVRs are found throughout the protein sequence, while 8 DVRs can be identified within the TprD_2_ sequences (Fig. 1). To obtain predictions of TprC and TprD_2_ structures from their amino acid sequences (Fig. 1) and map the DVRs on these models, we used the recently developed AlphaFold2 software (https://AlphaFold.ebi.ac.uk/) (37). These new models revealed remarkably similar structures for TprC and TprD/D_2_ and identified these proteins as relatively large β-barrel integral OMPs of 20 β-strands connected by ten external loops (ExLs, protruding toward the extracellular milieu), and nine periplasmic loops (Fig.2A). The local model quality, indicated by the pLDDT gradient was high in the transmembrane and periplasmic loop regions, and slightly lower in the predicted ExLs, suggesting conformational flexibility (Fig. 2B). Except for two substitutions (aa 407 and 410 mapping to a periplasmic β-turn; Fig.1), all DVRs localized within a subset of the surface-exposed ExLs (Fig.2C). More specifically, DVRs were located in ExL1, ExL5-6 and ExL8-10 of the TprC and TprD/D_2_ models; while ExL2-4 harbored conserved loops. ExL7 is also conserved between TprD_2_ sequences from various isolates, although its shows only 60% of sequence identity to the ExL7 of other TprC and TprD variants (Fig.1). *In silico* prediction of B-cell epitopes using BepiPred2.0 (http://www.cbs.dtu.dk/services/BepiPred/), IEDB (https://www.iedb.org/), and BCpreds (http://ailab-projects1.ist.psu.edu:8080/bcpred /data.html) (File S1) showed that the putative TprC and TprD ExLs were also enriched in immunogenic epitopes. Therefore, it is possible that the antigenic variability in the ExLs regions has functional significance in immunity to the *T. pallidum* subspecies. To validate the B-cell epitope prediction and evaluate the cross-reactivity of these epitopes across species and strains, we performed experimental B-cell epitope mapping of the TprC, and TprD/D_2_ proteins with a Geysan pepscan approach based on overlapping synthetic peptides (38) using sera from animals infected with *Tp* subsp. *pallidum*, *Tp* subsp. *pertenue*, and *Tp* subsp. *endemicum* strains. Furthermore, we compared antibody reactivity in sera from infected rabbits with that of sera from rabbits immunized with a subset of full-length refolded recombinant TprC proteins.

**Figure 1.**
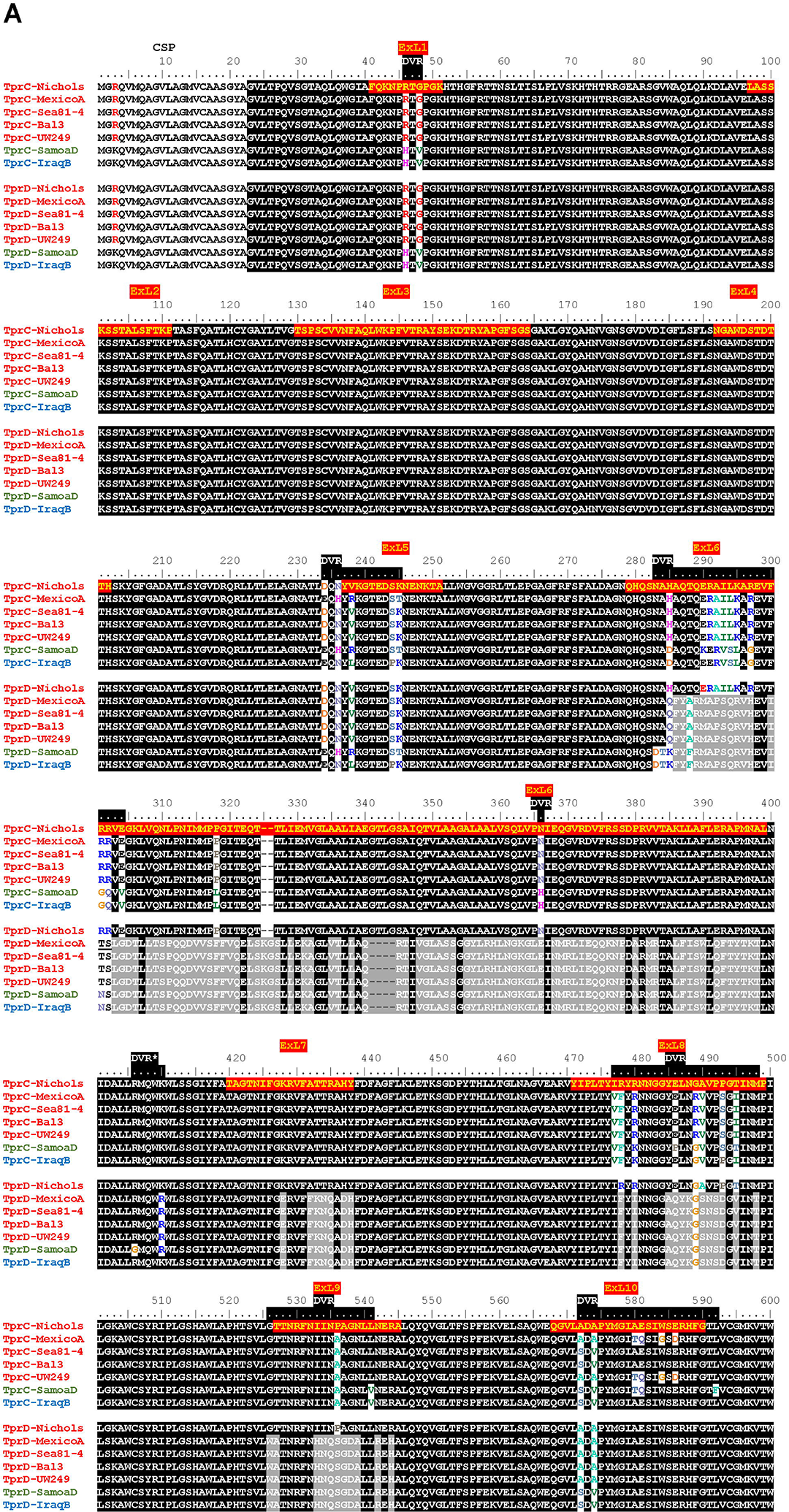
Alignment of amino acid sequences of the TprC and TprD/D_2_ variants. *Tp.* subsp. *pallidum* strains (Nichols, MexicoA, Sea81-4, Bal3, and UW249) are indicated in red font on the left of the sequence. The *Tp* subsp. *pertenue* strain (SamoaD) is in green font, and the *Tp* subsp. *endemicum* (IraqB) strain is in blue font. The Chicago and Bal73-1 TprC and TprD sequences (not shown) are identical to the Nichols strain. The MexicoA, Sea8-14, Bal3, UW249, SamoaD, and IraqB strains harbor a TprD_2_ variant within the *tprD* locus. CSP: predicted cleavable signal peptide; ExL: External Loop. Amino acids encompassing the ExLs predicted by AlphaFold2 are highlighted in red with yellow text only in the top sequence. DVR: Discrete Variable Region. DVRs are highlighted in black along the ruler. *Indicates a DVR found in TprD_2_ but not TprC and TprD variants.

**Figure 2.**
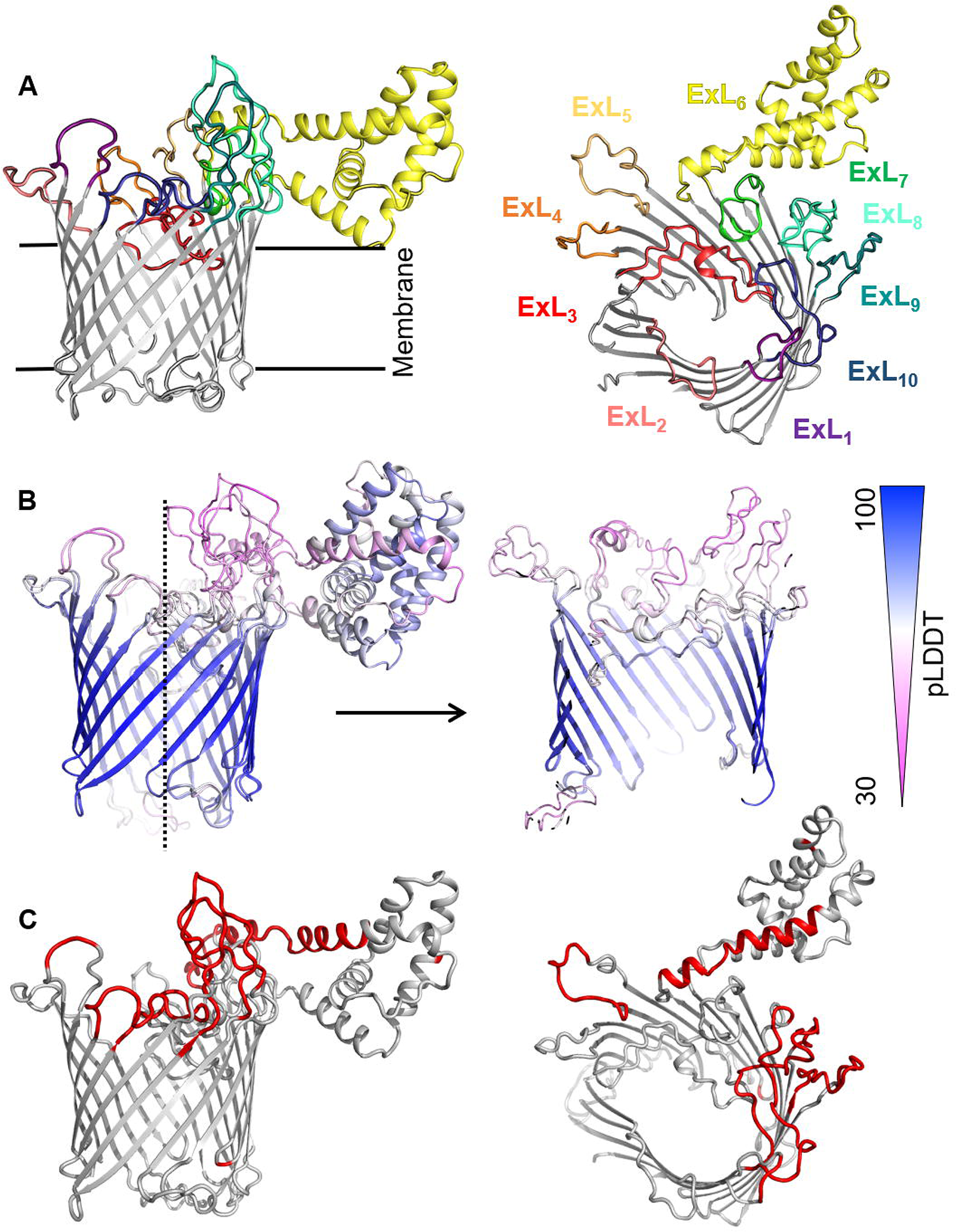
Predicted structures of TprC and TprD/D_2_ using AlphaFold2. **(A)** AlphaFold2 predicts very similar 20-strands β-barrel structures for both TprC and TprD/D2 proteins. TprC is shown on the figure and the 10 putative extracellular loops (ExLs) are color-coded. The model does not include the cleavable signal peptide (CSP; aa 1-22 in Fig.1). Once the predicted structures were superimposed to every available PDB structure by the DALI software (63) to identify structurally similar porins. DALI analysis results and PDB matches are reported in File S1. The highest-scoring structures did not the exact number of β-strands predicted by AlphaFold2 for TprC and TprD_2_ β-barrels, but slightly higher or slightly lower, but well within the models of integral OMPs with no large periplasmic domains. These results suggest that these Tpr proteins belong to a new family of porins not yet represented in the PDB. **(B)** The predicted structures of TprC and TprD/D_2_ are nearly identical in the transmembrane region (backbone root-mean-square deviation, or RMSD, = 0.75) where the estimated per-residue model confidence is very high (Predicted Local Distance Difference Test, pLDDT > 75). Full pLDDT analysis is reported in File S3 for TprC and TprD_2_. More differences are seen in the ExL regions, which also show lower pLDDT, suggesting structural flexibility of these loops. **(C)** The DVRs (colored in red) identified by aligning TprC and TprD_2_ sequences (Fig.1) of different strains localize in the predicted ExL regions.

### Humoral responses to homologous TprC and TprD/D_2_ peptides in experimentally infected rabbits

Serum samples from groups of three laboratory rabbits infected intratesticularly (IT) with one of seven syphilis strains (Nichols, Chicago, Bal73-1, MexicoA, Sea81-4, Bal3, and UW249), one yaws strain (SamoaD), and one bejel strain (IraqB) were obtained at day 30, 60, and 90 post-infection. Pooled sera from animals in each infection group/time point were tested in ELISA to assess reactivity to homologous overlapping synthetic peptides (20-mers overlapping by 10 amino acids) representing the TprC and TprD/D_2_ variants previously identified in each strain. The full list of synthetic peptides used in this study, with amino acids encompassing predicted ExLs highlighted in red with yellow text, and percentage amino acid homology among peptides across strains is shown in Table 1. Peptide nomenclature is explained in Table 1 footnote. Cumulative absorbance data from the three timepoints (sum of the mean absorbance values for day 30, 60, 90 values for each infected rabbit group) are reported in Fig.3A-C. Epitope mapping studies of the NH_2_-terminal portion of the protein resulted in the identification of six highly reactive peptide regions (Fig.3A) representing sequences shared by all TprC and TprD genes in the studied subspecies *pallidum* strains: C1-C3, C6, C13-C14, C18, C20, and C25-C29. Based on AlphaFold2 structural predictions, 9 of these 13 peptides had at least 7 amino acids mapping to the predicted external loops of the protein, while only four reside in predicted transmembrane transmembrane scaffolding and periplasmic loop regions (C1, C6, C20 and C25; Fig.1A, and Table 2). It is noteworthy that all three B cell epitope prediction programs uniformly predicted all six of the experimentally determined epitope-containing regions of the NH_2_-terminal portion of the subspecies *pallidum* TprC and D proteins (File S1).

**Figure 3.**
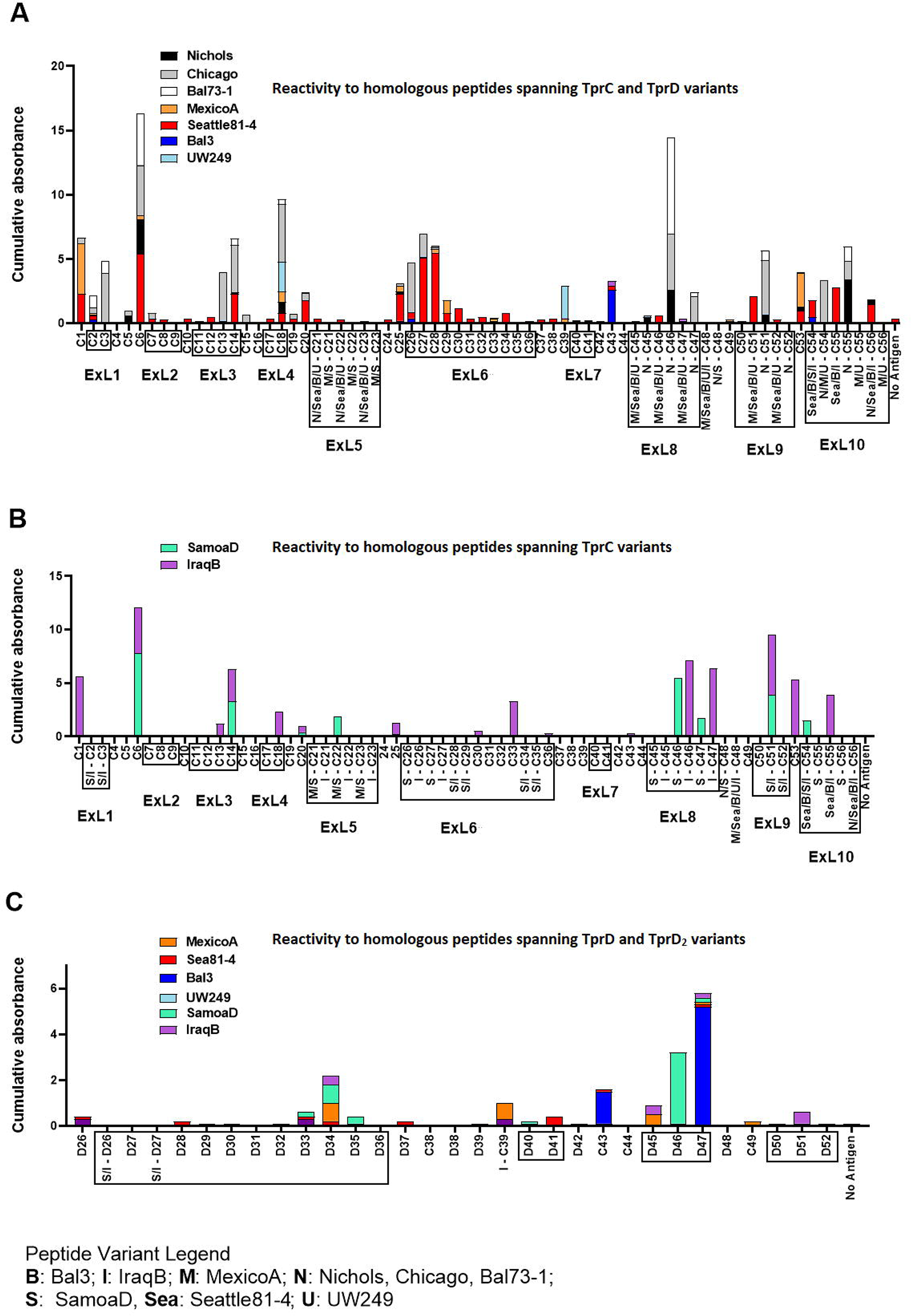
Reactivity of sera from experimentally infected animals to homologous peptides representing the TprC, TprD and TprD_2_ variants. (A) Reactivity to homologous peptides spanning TprC and TprD proteins of sera from rabbits infected with *Tp* subsp. *pallidum* (Nichols, Chicago, Bal73-1, MexicoA, Sea81-4, Bal3, and UW249B) collected at day 30, 60, and 90 post-infection. Nichols, Chicago, and Bal73-1 sequences are identical. (B) Reactivity to homologous peptides spanning TprC variants of immune sera from groups of rabbits infected with *Tp* subsp. *pertenue* (SamoaD) or *Tp* subsp. *endemicum* (IraqB) strains collected at day 30, 60, and 90 post-infection. (C) Reactivity to homologous peptides spanning TprD and TprD_2_ variants of sera collected at day 30, 60, and 90 post-infection from all TprD_2_-containing *Tp* subspecies and strains studied here. Cumulative Absorbance values are the sum of the mean OD values obtained from all animals in the infection group at all three time points. Boxed peptides contain at least seven amino acids (35% of the peptide length) belonging to a predicted ExL. Strain names on x axis are abbreviated as follows: N: Nichols; M: MexicoA; Sea: Sea81-4; B: Bal3; U: UW249; S: SamoaD; I: IraqB.

**Table 1.**
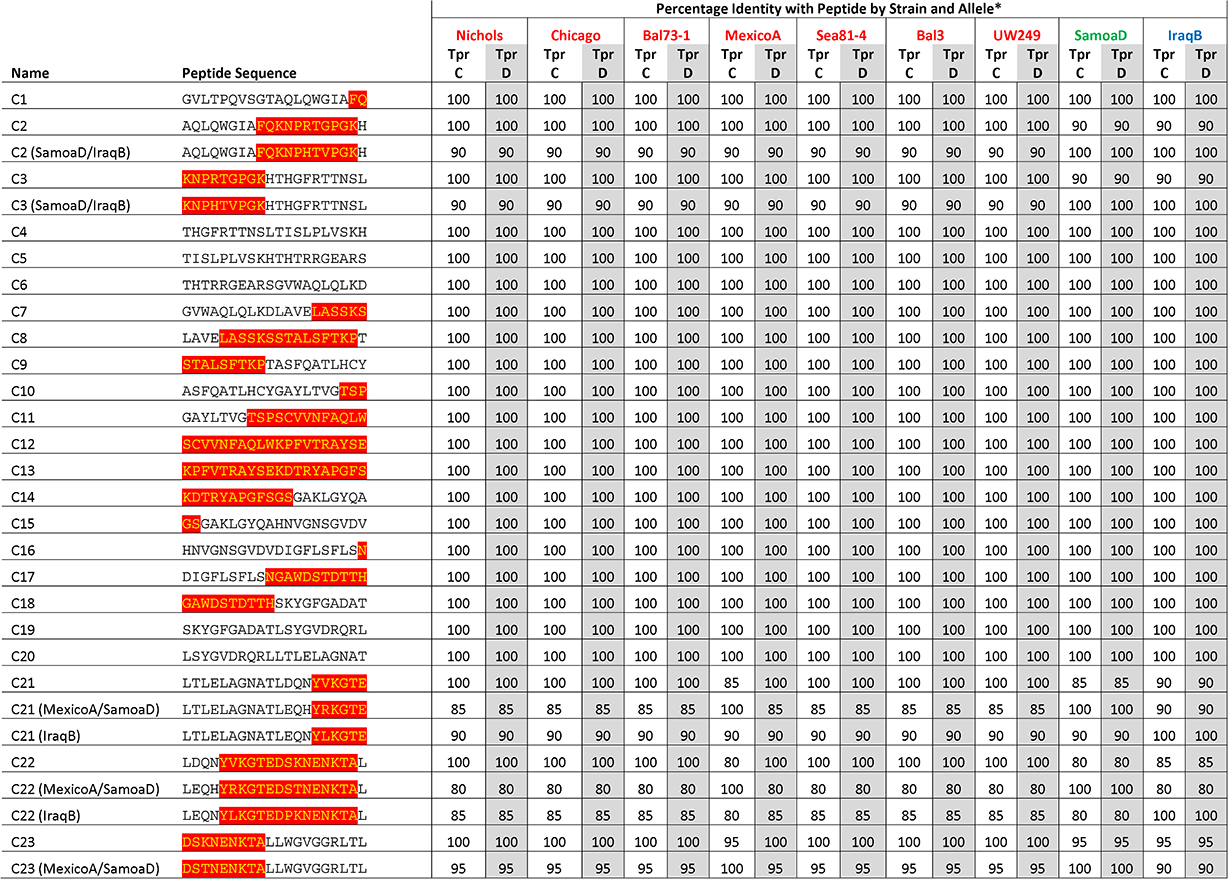
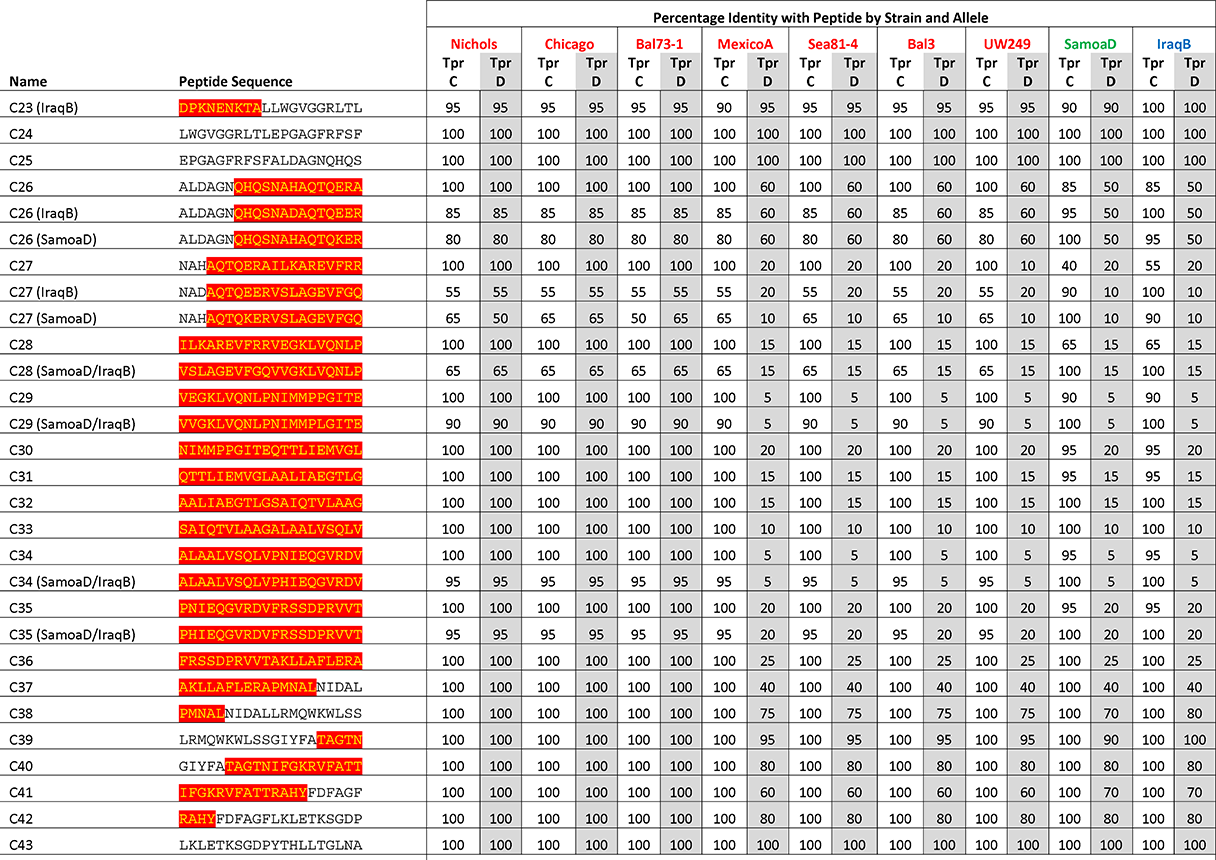
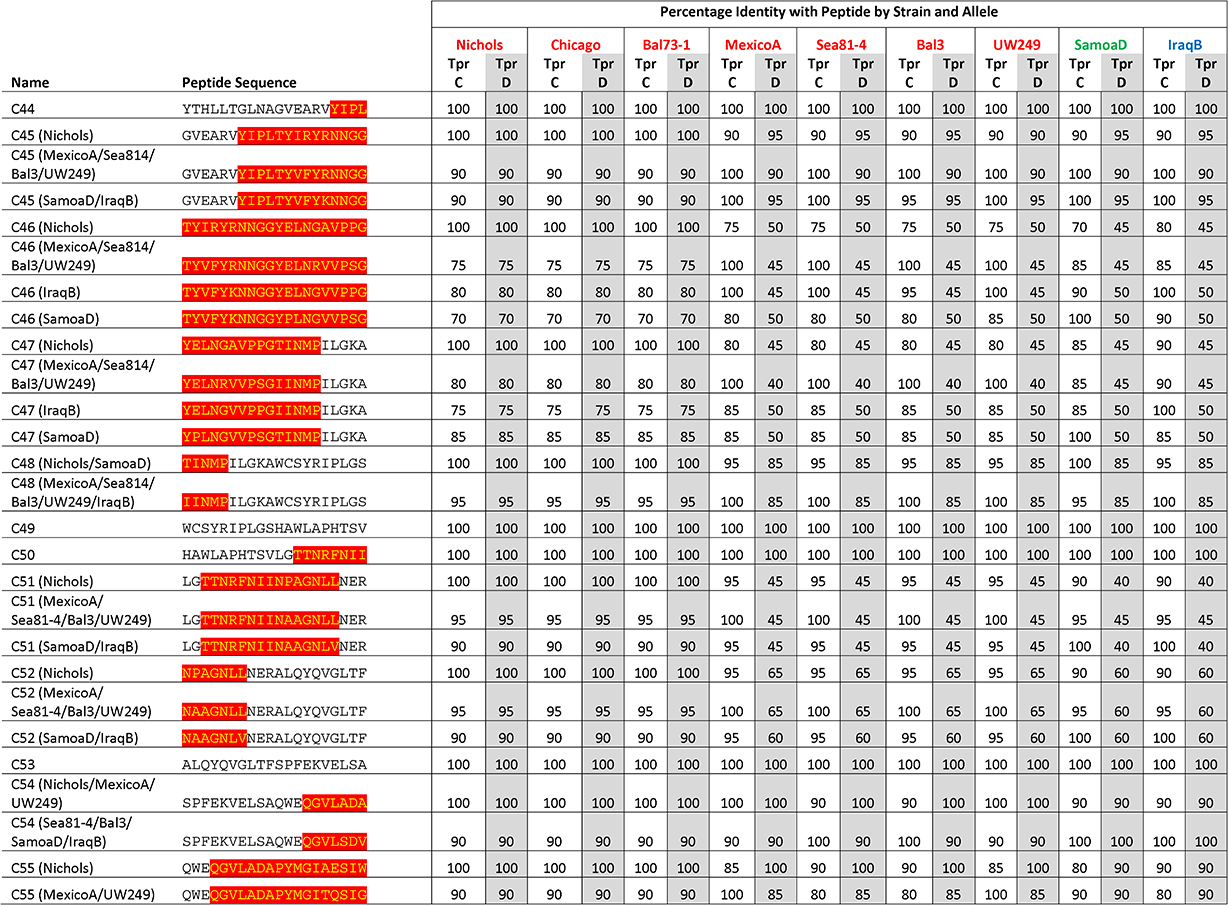
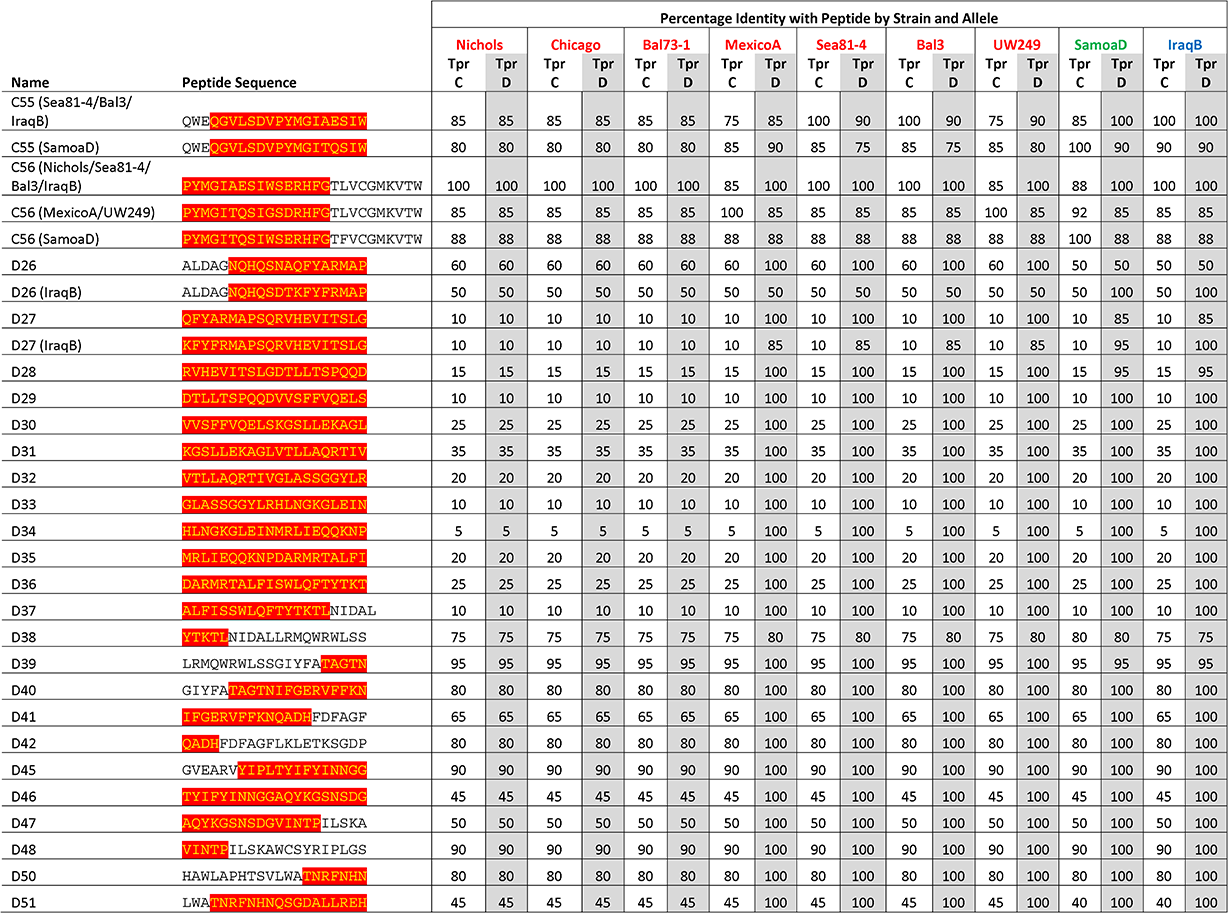
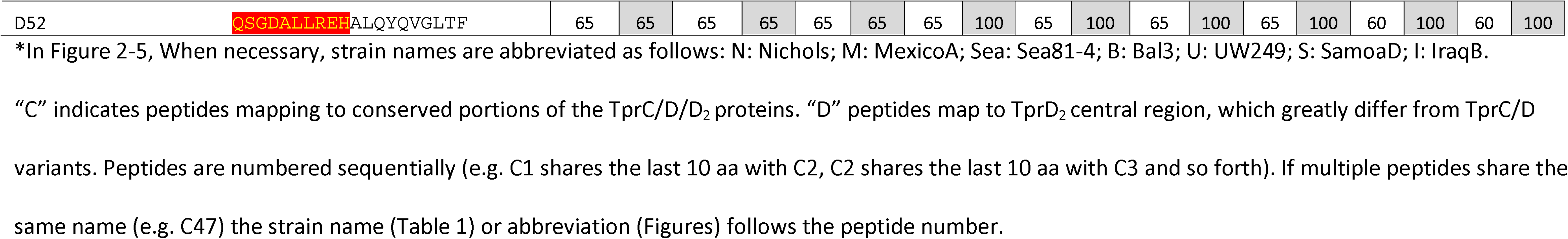
Synthetic peptides used in this study. Amino acids encompassing ExLs (predicted by AlphaFold) are highlighted in red with yellow text.

**Table 2.** Sequence of reactive peptides identified by infected-rabbit sera

| Sequences of reactive peptides based on Fig.2A (subspecies *pallidum*) |  |  |  |  |  |
| --- | --- | --- | --- | --- | --- |
| Peptide or peptide range | Experimentally determined Epitope-containing sequence | Location per AlphaFold | B-cell epitope predicted by |  |  |
|  |  |  | IEDB | BCpreds | BepiPred2.0 |
| C1-C3 | GVLTPQVSGTAQLQWGIAFQ<br>KNPRTGPGKHTHGFRRTNSL | Scaffold (C1)<br>and ExL1 (C2-3) | X | X | X |
| C6 | THTRRGEARSGVWAQLQLKD | Scaffold | X | X | X |
| C13-C14 | KPFVTRAYSEKDRYAPGFSGSGAKLGYQA | ExL3 | X | X |  |
| C18 | GAWDSTDTTHSKYGFGADAT | ExL4 | X | X | X |
| C20 | LSYGVDRQRLLTLELAGNAT | Scaffold | X |  |  |
| C25-C29 | EPGAGFRFSFALDAGNQHQ<br>NAHAQTQERAILKAREVFR<br>VEGKLVQNLPNIMMPPGITE | Scaffold (C25)<br>and ExL6 (C26-C29) | X | X | X |
| C39 | LRMQWKWLSSGIYFATAGTN | Scaffold | None |  |  |
| C43 | LKLETSGDPYTHLLTGLNA | Scaffold | X | X | X |
| N-C46-47 | TYIRYRNNGGYELNGAVPPGTINMPILGKA | ExL8 | X |  |  |
| C50-C51 | HAWLAPHTSVLGTTRNFNIINAAGNLLNER | ExL9 | X | X | X |
| C53-C55 | ALQYQVGLTFSPFEKVELSA<br>QWEQGVLS/ADV/APYMGIAESIW (N) or<br>QWEQGVLSVDPYMGIAESIW (Sea81-4/Bal3)* | Scaffold (C53)<br>and ExL10<br>(C54-C55) | X | X | X |
| Sequences of reactive peptides based on Fig.2B (subspecies *pertenue* and *endemicum*) |  |  |  |  |  |
| C1 | GVLTPQVSGTAQLQWGIAFQ | Scaffold | None |  |  |
| C6 | THTRRGEARSGVWAQLQLKD | Scaffold |  | X | X |
| C13-C14 | KPFVTRAYSEKDRYAPGFSGSGAKLGYQA | ExL3 | X | X | X |
| C18 | GAWDSTDTTHSKYGFGADAT | ExL4 | X | X |  |
| C20 | LSYGVDRQRLLTLELAGNAT | Scaffold | X |  |  |
| S – C22 | LEQHYRKGTEGSTNENKTAL | ExL5 | X | X |  |
| C25 | EPGAGFRFSFALDAGNQHQ | Scaffold |  | X |  |
| C33 | SAIQTVLAAGALAAALVSQLV | ExL6 | X |  |  |
| C36 | FRSSDPRVVTAKLLAFLERA | ExL6 | None |  |  |
| C43 | LKLETSGDPYTHLLTGLNA | Scaffold | X | X |  |
| S – C46 | TYVFYKNNGGYPLNGVVPSPG | ExL8 | X | X | X |
| I – C46 | TYVFYKNNGGYELNGVPPG | ExL8 | X | X | X |
| S – C47 | YPLNGVVPSTINMPILGKA | ExL8 | X | X | X |
| I – C47 | YELNGVPPGIINMPILGKA | ExL8 | X | X | X |
| C51 | LGTTRNFNIINAAGNLLNER | ExL9 | X | X | X |
| C53 | ALQYQVGLTFSPFEKVELSA | Scaffold | None |  |  |
| S – C54 | SPFEKVELSAQWEQGVLSDV | ExL10 | X | X | X |
| I – C55 | QWEQGVLSVDPYMGIAESIW | ExL10 | X | X | X |
| Sequences of reactive peptides based on Fig.2C |  |  |  |  |  |
| D34-D36 | HLNGKGLEINMRLEQQKNP<br>DARMRTALFISWLQFTYTKT | ExL6 | X | X | X |
| I – C39 | LRMQWKWLSSGIYFATAGTN | Scaffold | None |  |  |
| C43 | LKLETKSGDPYTHLLTGLNA | Scaffold | X | X | X |
| D46-D47 | TYIFYINNGGAQYKGSNSDGVINTPILSKA | ExL8 | X | X | X |
| D49 | WCSYRIPLGSHAWLAPHTSV | Scaffold | None |  |  |
| D51 | LWATNRFNHNQSGDALLREH | ExL9 | X | X | X |

Lightly shaded peptides are in the NH<sub>2</sub>-terminal portion of the protein.
\*S and V in the Sea 81-4/Bal3 strain, A and A in Nichols

Several epitopes were also identified in the COOH-terminal region of these proteins, and corresponded to peptides the same regions in *pallidum* and non-*pallidum* subspecies: C46 and C47 homologs from Nichols (Fig.3A, Table 2), SamoaD (S-C46, S-C47; Fig.3B and Table 2) and IraqB (I-C46, I-C47; Fig.3B and Table 2), C51 homologs from Nichols (N-C51), SamoaD and iraqB (S/I-C51) (Fig.3A-B and Table 2); and C53-C55 homologs from Nichols, Bal3/Sea81-4 (Fig.3A and Table 2), and IraqB (Fig.3B and Table 2). Similarly, the C43, and D45-D47 (ExL8) peptides, mapping to the TprD_2_ COOH-terminus were found to contain B-cell epitope(s) (Fig.3C and Table 2). Additional TprD_2_ peptides found to be reactive were D33-D35 (ExL6), I-C39, D40-41 (ExL7), C49, and D51 (ExL9). In our 3D models of these proteins, all the reactive peptides in the COOH-terminus fall within predicted ExLs (Tables 1-2 and Fig.1), except for C43, most of I-C39 (75%), C49, and C53, which are predicted transmembrane transmembrane scaffolding sequences. Of these “scaffold epitopes”, only one (C43) was predicted by a B-cell prediction program (File S1).

The percentage of immune sera that showed reactivity to many of the peptides was variable. For example, peptides C3, and C13 were recognized by rabbits infected with 28% of the *Tp* subsp. *pallidum* strains; peptides C1, and C27 were recognized by rabbits infected with 42% of the strains; peptides C14 and C28 were recognized by rabbits infected with 57% of the strains; C6 was recognized by rabbits infected with 71% of the strains; and peptides C2 and C18 were recognized by rabbits infected with 85% of the *Tp* subsp. *pallidum* strains (Fig.1). Overall, based upon the AlphaFold2 models, these results show that humoral reactivity elicited to these Tpr antigens during experimental infection is directed primarily to predicted surface-exposed regions of the TprC/D and TprD_2_ proteins.

### Reactivity to non-homologous TprC and TprD_2_ peptides

Epitope mapping using short peptides based on the TprC and TprD/D_2_ sequences from multiple *Tp* strains and sera from infected animals also allowed us to investigate cross-reactivity to non-homologous peptides to determine the fine specificity of the antibody response to these antigens. Such analyses focused on peptides mapping to the proteins’ COOH-terminal regions, due to the higher sequence variability in this region, compared to the more conserved NH_2_- terminal region (Fig.1). Major variable regions include peptides C46 - C47 (mapping to the predicted ExL8), C51 (ExL9), and C55 (ExL10) (Fig.1 and Tables 1-2). Four distinct variants of each of the C46, C47, and C55 peptides, and three variants of C51, representing all sequences found in the strains studied here, were tested against all nine pools of immune sera obtained at day-90 post-experimental infection.

As shown in Fig.4A-D, very few sera were reactive only to their homologous peptide. For example, the Bal73-1and SamoaD antisera were primarily reactive only to their own C46 sequences (Fig.4A), although the Bal73-1 antiserum showed a very modest reactivity to the IraqB peptide variant (Fig.4A). In contrast, none of the sera tested against the C47 variants exhibited reactivity against the homologous peptide, but the Chicago, Bal73-1, IraqB and Sea81- 4 sera were reactive against some heterologous variants (Fig.4B). Only Chicago and Bal73-1 sera showed complete strain-specificity for the C51 peptides (Fig.4C), while none of the sera reactive to C55 showed complete strain-specificity (Fig.4D). When tested against TprD2 peptides, most antisera did not show any reactivity. There were, however, two exceptions, as the Chicago sera cumulatively showed reactivity to the D34 and D47 peptides, with OD values of 3.6 and 6.6, respectively. However, only the D47 peptide was consistently recognized at all time points, while D34 was recognized only at day 60. Overall, these data indicate a relatively high level of cross immunity and perhaps suggest that immunization with a given sequence might generate cross-reactive antibodies able to overcome the obstacle of sequence diversity among TprC epitopes, a feature that is desirable in vaccine development as they may be broadly opsonic or neutralizing.

**Figure 4.**
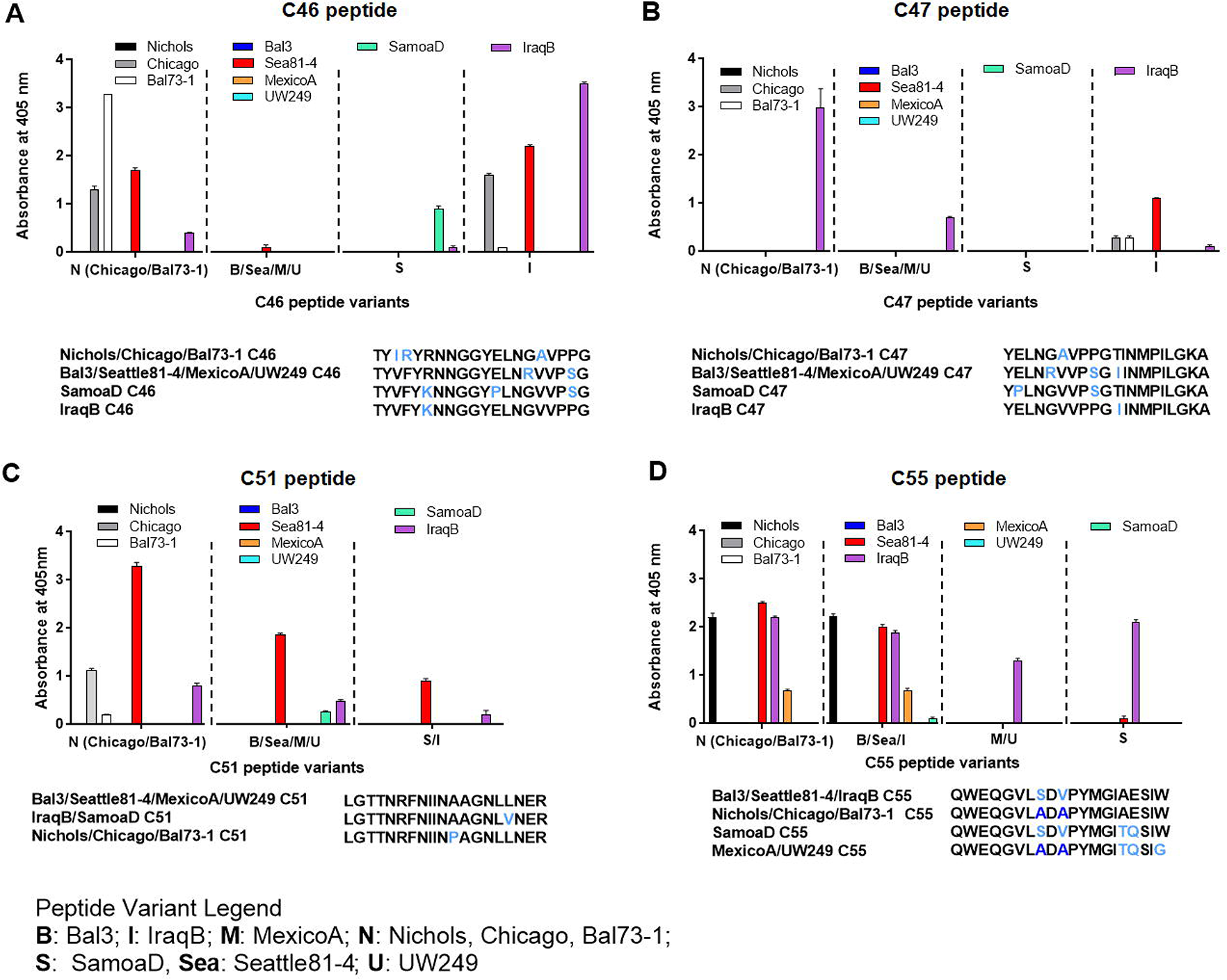
Reactivity of sera from experimentally infected animals to homologous and non-homologous peptides C46, C47, C51, and C55. Humoral reactivity of day-90 sera from experimentally infected animals to homologous and non-homologous TprC peptides. **(A-D)** reactivity to C46, C47, C51, and C55 variants. Strain names on x axes are abbreviated as follows: N: Nichols; M: MexicoA; Sea: Sea81-4; B: Bal3; U: UW249; S: SamoaD; I: IraqB.

### Humoral response to TprC peptides following rabbit immunization with full-length refolded antigens

Refolded antigens, analyzed using circular dichroism (CD), were found to have a β-barrel component of about 48% for all three antigen variants. Random coil was also found to be 48% of the protein structure, while only 4% was identified as alpha helices. Epitopes recognized following immunization with any of three recombinant full-length TprC variants from *Tp* subsp. *pallidum* strains (Nichols/Chicago/Bal73-1, Sea81-4/Bal3, and MexicoA) were also identified to evaluate differences with infection-induced immunity. Results showed that sera from animals immunized with the Nichols TprC sequence were highly reactive to peptides C1-C3, C6 and C47, and moderately reactive to peptides C5, C9, C16-18, C28, C32, C34-C35, C53 and C55 (Fig.5A). Of these 16 peptides, six mapped almost exclusively to putative surface-exposed loop regions (C28, C32, C34, C35, C47, and C55), five (C1, C5-C6, C16, and C53) mapped to predicted transmembrane transmembrane scaffolding sequences, while five peptides (C2-C3, C9 and C17-C18) contained both surface-exposed loops and scaffold regions. Sequences of these peptides and location in the predicted protein models are reported in Table 3. When tested against non-homologous peptides (Fig.5B), the Nichols TprC-immunized sera strongly recognized the SamoaD/IraqB C2-C3 variants, and all three heterologous C47 variants (SamoaD, Iraq B, and Sea81-4), while modest reactivity was seen towards the MexicoA/UW249 C55 peptide variant, the SamoaD/IraqB C34, and both C26 variants from SamoaD and IraqB (Fig.5B). Immunization with the Bal3 variant of TprC elicited high reactivity to peptides C1-3, C6, and C13, and moderate reactivity to peptides C7, C16-18, C20, C43, C47, and C49 (Fig.5C, Table 3). Of these thirteen peptides, six (C2-C3, C13, C17-C18, and C47) mapped predominantly to ExLs, and seven (C1, C6, C7, C16, C20, C43, and C49) predominantly to the protein transmembrane transmembrane scaffolding (Table 3). Cross-reactivity to non-homologous peptides was seen predominantly to the SamoaD/IraqB C2 and C3, IraqB C22, and all variants of C47 and C51 (Fig.5D). Antisera from rabbits immunized with the MexicoA TprC variants primarily recognized homologous peptides C1-3 and, secondarily, C5, C6, and C28 (Fig.5E). Of these six, one peptide mapped to the predicted ExL6 (C28), three mapped only to the transmembrane transmembrane scaffolding (C1, C5-C6) and two mapped to a peptide predicted to contain portions of both (C2-C3) (Table 3). Cross-reactivity to the non-homologous SamoaD/IraqB C2 and C3 was also detected (Fig.5F). Overall, these data show that, as seen in infection-induced immunity, the humoral response following immunization with full-length TprC variants is mainly elicited by predicted surface-exposed sequences, rather than sequences mapping to the β-barrel transmembrane scaffolding, and that cross-reactivity to non-homologous peptides is common.

**Figure 5.**
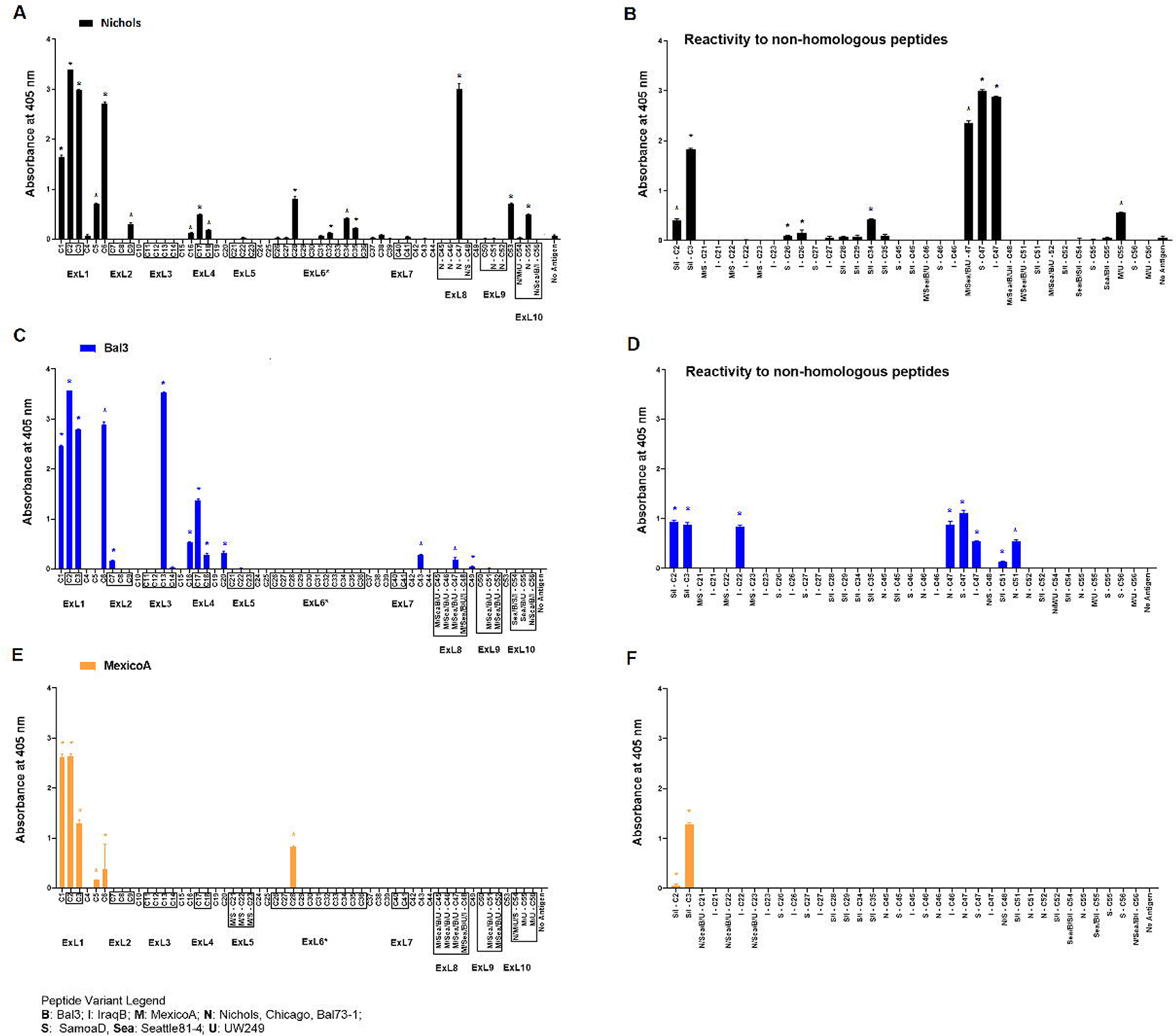
Humoral reactivity to TprC peptides following immunization with refolded recombinant full-length TprC antigens. Reactivity to TprC homologous (left panels) and non-homologous peptides (right panels) in sera from rabbits immunized with Nichols **(A, B)**, Bal3 **(C, D)**, and Mexico A **(E, F)** variants of TprC. Asterisk (*) indicates significant reactivity compared to no antigen control. Peptides encompassing sequences predicted to be within ExLs are boxed. Peptide sequence and homology among strains are reported in Table 1. Strain names on x axes are abbreviated as follows: N: Nichols; M: MexicoA; Sea: Sea81-4; B: Bal3; U: UW249; S: SamoaD; I: IraqB.

**Table 3.** Sequences of reactive peptides following rabbit immunization

| Sequences of reactive peptides based on results shown in Fig.4A.<br>Immunization with TprC full-length Nichols variant |  |  |
| --- | --- | --- |
| Peptide or peptide range | Experimentally determined Epitope-containing sequences | Location per AlphaFold |
| C1-C3 | GVLT PQVSGTAQLQWGIAFQ<br>KNPRTGPGKHTHGFRTTNSL | Scaffold (C1) and ExL1 (C2-C3) |
| C5-C6 | TISLPLVSKHHTHRRGEARS<br>THTRRGEARSGVWAQLQLKD | Scaffold |
| **C9<sup>1</sup>** | **STALSFTKPTASFQATLHCY** | ExL2/Scaffold |
| C16-18 | HNVGNSGVDVDIGFLSFLSN<br>GAWDSTDTTHSKYFGADAT | Scaffold (c16-17)/ExL4 (C17-18)/Scaffold (C18) |
| C28 | ILKAREVFRRVEGKLVQNLP | ExL6 |
| C32 | AALIAEGTLGSAIQTVLAAG | ExL6 |
| C35 | PNIEQGVRDVFRRSDPRVVT | ExL6 |
| N/Sea/U - C47 | YELNGAVPPGTINMPILGKA | ExL8 |
| C53 | ALQYQVGLTFSPFEKVELSA | Scaffold |
| C55 | QWEQGVLSDPYMGIAESIW | ExL10 |
| Sequences of reactive peptides based on results shown in Fig.4B.<br>Immunization with TprC full-length Bal3 variant |  |  |
| C1-C3 | GVLT PQVSGTAQLQWGIAFQ<br>KNPRTGPGKHTHGFRTTNSL | Scaffold (C1) and ExL1 (C2-C3) |
| C6-C7 | THTRRGEARSGVWAQLQLKDLAVELASSKS | Scaffold (C6) and ExL2 (C7) |
| C13-C14 | KPFVTRAYSEKDTRYAPGFSGSGAKLGYQA | ExL3 |
| **C16<sup>1</sup>-C18** | **HNVGNSGVDVDIGFLSFLSN**<br>GAWDSTDTTHSKYFGADAT | Scaffold (C16) and ExL4 (C17-C18) |
| C20 | LSYGVDQRLLTLELAGNAT | Scaffold |
| C43 | LKLETKSGDPYTHLLTGLNA | Scaffold |
| C47 | YELNRVPSGIINMPILGKA | ExL8 |
| C49 | WCSYRIPLGSHAWLAPHTSV | Scaffold |
| C51 | LGTTNRFNIINAAGNLLNER | ExL9 |
| Sequences of reactive peptides based on results shown in Fig.4C.<br>Immunization with TprC full-length MexicoA variant |  |  |
| C1-C3 | GVLT PQVSGTAQLQWGIAFQ<br>KNPRTGPGKHTHGFRTTNSL | Scaffold (C1) and ExL1 (C2-C3) |
| C5-C6 | TISLPLVSKHHTHRRGEARS<br>THTRRGEARSGVWAQLQLKD | Scaffold |
| **C9<sup>1</sup>** | **STALSFTKPTASFQATLHCY** | Scaffold/ExL2 |
| C28 | ILKAREVFRRVEGKLVQNLP | ExL6 |

<sup>1</sup> C9/C16 peptides were not predicted to harbor B-cell epitopes by IEDB, BCpreds and bepiPred2.0

A side-by-side comparison of the infection- vs. immunization-induced humoral response to peptides is shown in Fig.6. For this comparison, the mean value of the cumulative reactivity seen in sera at day 30, 60, and 90 sera post-experimental infection is shown for each peptide. Sera from immunized animals were obtained three weeks after the last immunization. All sera were tested at the same dilution. In general, immunization-induced reactivity to most peptides appeared to be higher than that elicited by experimental infection; specific examples include C1-C3, C5, C9, C16-C17, C28, C32, C34-C35, N-C47, and C53 peptides (Fig.6A). For Nichols-clade *T. pallidum* strains (Fig 6A), which contain identical *tprC* and *tprD* loci, this was most noticeable for epitopes located in the NH_2_- and COOH-terminal regions of the protein. In contrast, infection-induced antibody responses to epitopes in the central part of the protein were comparable to or higher than those induced by immunization. For TprD2-containing subsp. *pallidum* strains (Fig.6B and 6C), immunization-induced responses were limited to the NH_2_- terminal portion of the protein (including ExL1-3) and virtually no immunization-induced antibodies were detected for epitopes in the central and COOH-terminal regions, although these were recognized by infection-induced responses. Overall, these data support that, in most cases, immunization elicits a higher reactivity to TprC B-cell epitopes compared to experimental infection, particularly for those epitopes located in the NH_2_-terminal portion of the protein. These data support the preferential use of the amino portion of TprC, which contains multiple conserved ExLs, for vaccine studies.

**Figure 6.**
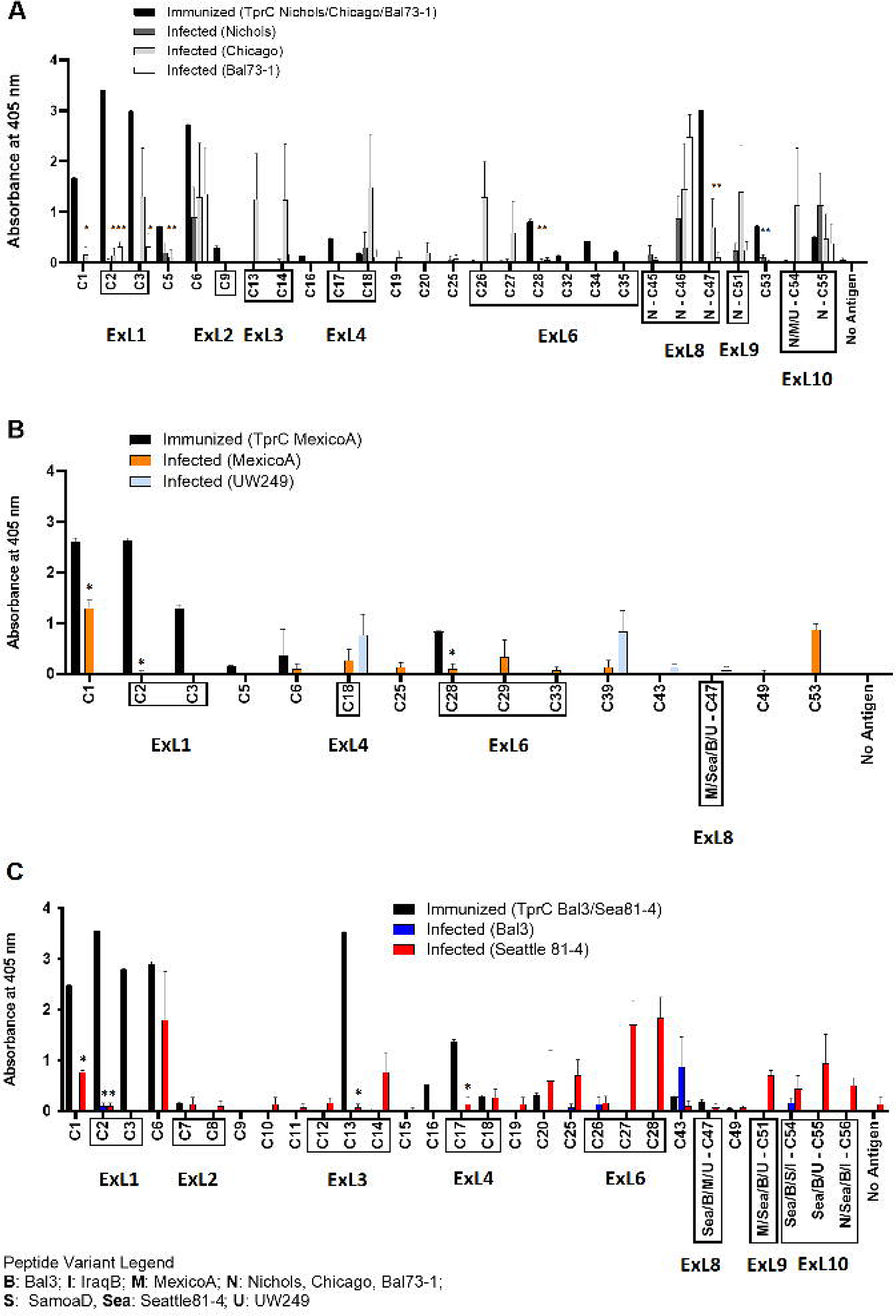
Comparison of reactivity of sera from infected animals vs. immunized animals. **(A-C)** Reactivity to peptides following immunization with TprC variants compared to experimental infection. Data shown are means +/- SEM of 3 rabbits per group: 3 weeks post final boost (immunized) and mean +/- SEM of values for days 30, 60, 90 post-infection (infected). Asterisk (*) indicates a significant difference in reactivity compared to the reactivity value following immunization. Peptides encompassing sequences predicted to be within ExLs are boxed. Strain names on x axes are abbreviated as follows: N: Nichols; M: MexicoA; Sea: Sea81-4; B: Bal3; U: UW249; S: SamoaD; I: IraqB.

## DISCUSSION

The continuing prevalence of syphilis, in the face of highly effective therapy and active control programs, highlights the need for a protective vaccine. The development of such a vaccine calls for a deeper understanding of the mechanisms of protective immunity and the antigens and adjuvants that induce protection. Our laboratories have been examining these issues for many years (22, 31, 39–47). Much of that work has focused on the Tpr antigens of *T. pallidum*. In this current study, B-cell epitope mapping studies of the TprC/D and TprD_2_ proteins of *Tp* reveal that antibodies arising during experimental infection recognize sequences predicted, using state-of-the-art modeling systems, to fall largely in the proteins’ surface-exposed loops. Because opsonic antibodies are required for efficient ingestion and killing of *T. pallidum* by macrophages, surface-exposed epitopes are attractive targets as vaccine candidate antigens.

A broadly protective vaccine would need to be effective against most strains of *T. pallidum*, optimally including the agents of syphilis as well as the endemic treponematoses yaws and bejel. Because some of the external loops of Tpr C/D and TprD_2_ demonstrate sequence heterogeneity among strains and subspecies of *T. pallidum*, we expected that these epitopes might be strain-specific, similar to the specificity demonstrated for the variable regions of TprK (41, 43, 48). For this reason, we included seven strains of *Tp* subsp. *pallidum* as well as strains from the subspecies *pertenue* and *endemicum* in our work. Unexpectedly, we saw considerable cross reactivity of antibodies toward the variant peptides (Fig.4). These findings support the use of TprC/D as at least one component of broadly effective candidate vaccine.

The AlphaFold2 structural predictions for TprC/D and TprD_2_, as well as our CD analyses of purified refolded recombinant TprC variants, support our model (35) that these Tprs are membrane-localized 20-stranded β-barrel proteins containing numerous surface-exposed loops. Very similar models for TprC were previously obtained using I-TASSER (49) (https://zhanggroup.org/I-TASSER/) (35). Interestingly, when AlphaFold2 and I-TASSER results are compared, the only difference is that I-TASSER splits ExL6 (Fig.1) into two external loops separated by a β-hairpin, so that I-TASSER predictions harbor 11 external loops instead of 10. AlphaFold2, on the contrary, predicts a significantly larger ExL6, mapping approximately to the proteins’ central domains. AlphaFold2 is the new standard for *ab-initio* structural prediction, and in the 2020 Critical Assessment of protein Structure Prediction (CASP) global challenge, it outperformed any other structure prediction algorithm, including I-TASSER (https://predictioncenter.org/casp14/zscores_final.cgi ). Furthermore, in a recent preprint (50), AlphaFold2 was shown to work well on structural prediction for membrane proteins, although the exercise focused mostly on alpha-helical membrane proteins, and additional analyses are necessary to establish the same benchmark for β-barrel proteins.

In previous work by Anand *et al.* (34, 51) significantly different models for the TprC/D proteins were reported, compared to those provided here. These models, however, are not supported by AlphaFold2, which finds the structure of all Subfamily I and Subfamily II Tpr family members to be very similar to the structures for TprC/D and TprD_2_ in Fig.2A. Although there is not unanimous agreement on the structure of these antigens within our scientific community, our epitope mapping data support our AlphaFold2 models, predicting a predominantly β-barrel structure for TprC and TprD/D2 (34, 51). Further studies and integration of all the structural, functional, and immunological data are needed to establish a consensus on the structure of these antigens until crystallographic (or equally reliable) data become available.

This study also provides evidence that infection with different strains might lead to differences in the breadth and intensity of the humoral response against the same epitope, as reported previously for responses to longer portions of the Tpr proteins (52). It is the case, for example, of rabbits infected with the Sea81-4 strain of *Tp* that overall recognize more TprC/D peptides compared to other *Tp* subsp. *pallidum* strains. The biological basis for these differences is unclear at this time, in part due to the limitations of our understanding of *Tp* biology and syphilis pathogenesis. As the technical gap in the approaches to study this difficult pathogen narrows, and genomics, proteomics, and transcriptomics data populate public repositories, more light will be shed on the causes of differential reactivity. Overall, however, it is plausible to postulate that enhanced serological reactivity might be due to an overall increased expression of the target antigen in each strain. This hypothesis is supported by previous work where we showed the *tprC* mRNA level was higher in the Sea81-4 strain compared to other *Tp* subsp. *pallidum* strains (Nichols, Chicago, Bal73-1) used in this study (53).

Our studies further demonstrated that epitopes in TprC/D and TprD_2_ are nearly-uniformly distributed across the length of the protein, even though the most reactive peptide epitopes are in the NH_2_- and COOH-terminal regions. Previously published (47) and ongoing experiments have shown that both of these regions in the Nichols TprC protein contain protective epitopes, as immunization with these antigen fragments significantly attenuated lesion development upon infectious challenge (47), and polyclonal antisera elicited by immunization with these portions facilitated treponemal ingestion by macrophages in opsonophagocytosis assays compared to normal rabbit sera (47) . Further work, however, will be necessary to identify which specific surface-exposed sequences provide targets for opsonic antibodies, which may not coincide with sero-dominant epitopes, as the pathogen gains an obvious advantage by exposing to the immune system epitopes with little or no protective value.

Protective B-cell epitopes (contrary to T-cell epitopes) are often conformational, and even when a significant portion of an epitope appears to be a short linear peptide, as in our study, it does not necessarily mean that the peptide represents the full epitope or, if it does, that the sequence will not require a certain conformation to elicit optimal bioactivity. For this reason, in the immunization studies performed in this study, we used CD-confirmed refolded recombinant antigens. The immunization-induced antibodies generally identified the same epitopes seen in infection, supporting the role of refolding in mimicking native structure, but immunization also resulted in recognition of a broader range of epitopes than seen during infection, including transmembrane scaffolding regions. This is likely because the scaffold regions are not shielded by the outer membrane in an immunization setting and are thus more easily processed for recognition. Thus, the design of vaccine immunogens is critical. Possible approaches vary from placing epitopes within chimeric antigens that could work as scaffold or, alternatively, using portions of the protein containing protective epitopes as structural elements of the antigen, or even using single β-hairpins instead of the full-length antigens. The work reported here represents an important step in evaluating TprC/D and TprD_2_ epitopes as part of the process that will lead to an effective vaccine for syphilis.

## MATERIALS AND METHODS

### Ethics Statement

New Zealand White rabbits were used for propagation of *T. pallidum* subspecies and strains and for experimental infections. Animal care was provided in accordance with the procedures described in the Guide for the Care and Use of Laboratory Animals (54) under protocols approved by the University of Washington Institutional Animal Care and Use Committee (IACUC, PI: Sheila Lukehart). The protocol number assigned by the IACUC committee that approved this study is 2090-08. No human samples were used in this study.

### Strain propagation and experimental infection

Outbred adult male New Zealand White rabbits ranging from 3.0-4.0 Kg were obtained from R&R Rabbitry (Stanwood, WA). Prior to entry into the study, serum from each animal was tested with both a treponemal (FTA-ABS) and a non-treponemal (VDRL; BD, Franklin Lakes, NJ) test to rule out infection with the rabbit syphilis agent *Treponema paraluiscuniculi*. Only rabbits seronegative in both tests were used for either propagation or experimental infection for sample collection. *Tp s*trains were propagated by intratesticular (IT) inoculation and harvested at peak orchitis as previously described (55). For experimental infections, groups of three rabbits were infected IT with a total of 5×10^7^ *Tp* cells per testis. In total, nine *Tp* isolates (one isolate per rabbit group) were used: seven *Tp* subsp. *pallidum* isolates (Nichols, Chicago, Bal73- 1, Sea81-4, Bal3, MexicoA, and UW249), one *Tp* subsp. *endemicum* (IraqB) and one *Tp* subsp. *pertenue* (SamoaD) (Table 4). Briefly, on the day of infection bacteria were extracted from rabbit testes in sterile saline containing 10% normal rabbit serum (NRS), and testicular extract was collected in sterile 15-ml tubes. Extracts were centrifuged twice at 1,000 rpm (180 x g) for 10 minutes in an Eppendorf 5810R centrifuge (Eppendorf, Hauppauge, NY) to remove gross rabbit cellular debris. Treponemes were enumerated under a dark-field microscope (DFM) and percentage of motile organisms was recorded. Extracts were then diluted in serum-saline to the desired concentration (5×10^7^/ml). Following IT injection, treponemal motility was assessed again to ensure that the time elapsed before injection into the new host did not affect pathogen viability. After IT inoculation, establishment of infection was assessed by monitoring development of orchitis during the following three weeks as well as by performing FTA-ABS and VDRL tests on sera collected at day 30 post-inoculation. Immune sera were collected from the animals at day 30, 60, and 90 post-infection. Animals were then euthanized. Extracted sera were heat-inactivated at 56°C for 30 min and stored at -20°C until use for ELISAs.

**Table 4.**
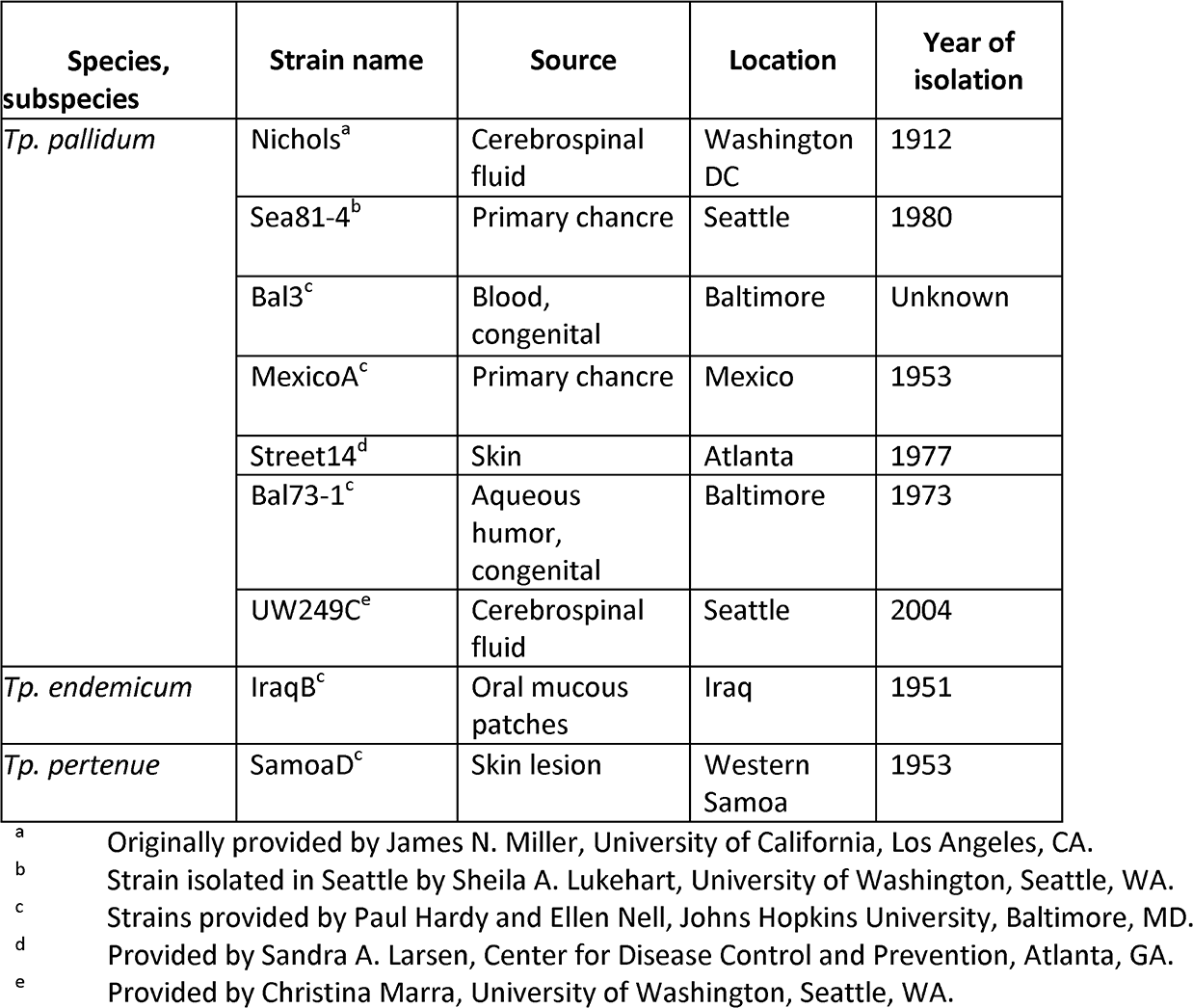
Treponemal strains used in this study

### Amplification and cloning of full-length *tprC* gene variants for expression of recombinant antigens

Sequences for the *tprC* gene of *Tp* isolates (Nichols, Sea81-4, and MexicoA) were previously cloned (35). For expression, the *tprC* sequences were sub-cloned into the pET23b+ vector (Life Technologies) between <u>BamHI</u> and <u>HindIII</u> using the primers C-S (5’-cg<u>ggatccg</u>atgg gcgtactcactccgca) and C-As (5’-gc<u>aagctt</u>ccatgtcactttcattccac). For sub-cloning, the *tprC* ORF was amplified in a 100-μl final volume using 0.4 units of GoTaq polymerase (Promega) with approximately 10 ng of DNA template. MgCl_2_ and dNTP final concentrations were 1.5 mM and 200 μM, respectively. Initial denaturation and final extension (72°C) were for 10 min each. Denaturation (94°C), annealing (60°C), and extension (72°C) were carried out for 1 min each for a total of 35 cycles. Amplicons were purified, digested, and ligated into the pET23b+ vector. As a result of cloning into pET23b+, 28 additional amino acids were added to the TprC ORFs (14 NH_2_- terminal and 14 COOH-terminal amino acids), including the COOH-terminal 6×His tag for affinity purification. Ligation products were used to transform OneShot TOP10 chemically competent *E. coli* cells (Life Technologies) according to the provided protocol. Transformations were plated on LB-Ampicillin (100 μg/ml) agar plates for selection. For each cloning reaction, individual colonies were screened for the presence of insert-containing plasmids using primers annealing upstream and downstream of the pET23b+ vector poly-linker (T7 promoter and terminator primers). Positive plasmids were extracted from overnight liquid cultures obtained from replica colonies by using the Plasmid Mini kit (Qiagen, Germantown, MD), and two to five clones for each strain were sequenced to ensure sequence fidelity to the previously cloned templates (35). For expression of recombinant antigens, a suitable clone for each *tprC* gene variant was used to transform *E. coli* Rosetta (DE3) competent cells (Life Technologies).

### Expression, purification and refolding of recombinant proteins

*E. coli* cells were grown overnight in LB media supplemented with ampicillin (100 μg/ml). The following day, multiple flasks containing 200 ml of auto-inducing media (56), were inoculated with 20 ml of overnight culture in a 2-liter baffled flask and grown at room temperature for 72 h at 175 rpm in a shaking incubator. Expression of recombinant antigens in induced and un-induced controls was assessed by immunoblot using a monoclonal anti-poly- histidine antibody (Millipore-Sigma, diluted 1:2000) after SDS-PAGE. Prior to purification, presence of the recombinant protein in the soluble and insoluble cellular fractions was evaluated by SDS-PAGE and immunoblot. Recombinant TprC purification was carried on under denaturing conditions. Briefly, *E. coli* cell pellets were resuspended in 5 ml/g of dry culture weight of binding buffer (50 mM NaH_2_PO_4_, 10 mM imidazole, pH 8.0) w/o denaturing agent, and the suspension was sonicated in ice with 100 pulses of 6 s each, with each pulse being separated by 10-s intervals. Insoluble components (containing the desired products) were precipitated by centrifugation and resuspended in 5 ml/ g of culture weight of binding buffer (50 mM NaH_2_PO_4_, 5 mM imidazole, pH 8.0) containing 6M Guanidine-HCl denaturing agent and sonicated again as above. Insoluble components were precipitated again by centrifugation and the supernates were saved. For affinity chromatography, 5.0 ml of nickel-agarose (Ni-NTA agarose, Qiagen) was packaged into a 1.5×14 cm column (Bio-Rad, Carlsbad, CA) and washed with 3 column volumes of molecular-grade H_2_O and 6 column volumes of binding buffer + denaturing agent. Cell lysate was then loaded, and the flow was adjusted to 1 ml/min. Unbound proteins were washed using 10 bed volumes of binding buffer, followed by 6 column volumes of wash buffer (50 mM NaH_2_PO_4_, 20 mM imidazole, pH 8.0) containing denaturing agent. Washing continued until the *A*_280_ of the flow through was <0.01 AU. Recombinant TprC was eluted with 15 ml of elution buffer (50 mM NaH_2_PO_4_, 300 mM imidazole, pH 8.0) containing denaturing agent. Eluted fractions devoid of visible contaminants by SDS-PAGE and Coomassie staining were pooled, and protein concentration was assessed by micro-bicinchoninic (BCA) assay (Thermo-Fisher). Pooled fractions were then dialyzed in PBS using a 10 kDa MWCO Slide-A-Lyzer dialysis cassette (Thermo-Fisher) over 12 hours, ensuring PBS change every ∼4 hours. Precipitated protein, resulting from elimination of Guanidine-HCl during dialysis was transferred into microcentrifuge tubes and spun down at full speed. After removing the supernate, the pellet was resuspended in a volume of PBS containing 6M urea suitable to achieve a protein concentration of ∼4 mg/ml, and protein concentration was then reassessed using the micro-BCA assay kit (Thermo-Fisher). Prior to immunizations, urea was eliminated using Profoldin (Hudson, MA) M7 renaturing columns for membrane proteins, which were used according to the manufacturer’s protocol. M7 renaturing columns were found to provide the best yield when screened along with 19 other conditions offered by Profoldin. Lipid composition of the elute buffer included lysophosphatidylcholine (∼5 mM), arginine (∼150 mM), glycerol (∼10%), dodecyl maltoside (0.7 mM), and Tris-HCl (0.1 mM), pH 7.5). Following buffer exchange, soluble protein concentration was evaluated using micro-BCA assay and analyzed by circular dichroism (CD) to evaluate percentage of β-sheet, alpha-helix, and random coil. CD spectra (190 to 260 nm) were acquired in triplicate at room temperature using 0.5 mg/ml of recombinant refolded TprC in a Jasco-1500 high-performance CD spectrometer. CD spectra were analyzed using the online platform Dichroweb (http://dichroweb.cryst.bbk.ac.uk/html/home.shtml ) (57) and the spectra from buffer alone for background subtraction.

### Rabbit immunization

Groups of three rabbits each were immunized with one of the purified, refolded recombinant TprC variants. Rabbits were injected with 125 μg of refolded protein every 3 weeks for a total of three immunizations. Prior to injection, antigen was mixed with an equal volume of in Titermax Gold Adjuvant (Millipore-Sigma), a water-in oil emulsion containing squalene, the block co-polymer CRL-8300, and a microparticle stabilizers to obtain a final volume of 1 ml. Immunogen-adjuvant preparation was performed according to the manufacturer’s instruction, and immunizations were performed via four 250 μl injections (each containing 31.25 μg of protein) into 4 intramuscular sites. Three weeks after the last boost, immunized animals were deeply anaesthetized, bled through cardiac puncture, and then euthanized.

### ELISA using synthetic peptides

Overlapping synthetic peptides (20-mers overlapping by 10 aa) were designed to represent the sequences of all TprC and TprD/D_2_ loci present in each of the seven strains examined in this study starting after the predicted signal peptide (AA 1-22; Fig.1 and Fig.2). Only the C56 peptide and its variants (Table 1), which represent the proteins’ COOH-terminus, were synthesized as 26-mers. A total of 120 peptides (Table 1) were produced by Genscript (Piscataway, NJ). Upon receipt, lyophilized peptides were rehydrated in sterile PBS to a stock solution of 200 μg/ml. Solubility of hydrophobic peptides was increased by adding up to 4% (v/v) DMSO per manufacturer’s instruction when needed (peptides C1, C4-7, C10, C15-16, C20, C25, C38-C39, C43-44, C53; Table 1). Reconstituted peptides were stored at -20°C until use. For ELISA, peptides were further diluted to 10 μg/ml in PBS, and 50 μl of working dilution (500 ng total) were used to coat the wells of a 96-well Microwell Maxisorp flat-bottom plate (Thermo-Fisher, Waltham, MA) as previously described (42). Absorbance was measured at OD_405_ using a Molecular Devices SpectraMax Plus microplate reader (Molecular Devices, San Jose, CA). A micro-BCA protein assay (Thermo Fisher) was performed in plates coated with Ag and washed to demonstrate that all peptides bound to the well surfaces in the plates (data not shown). For each serum from each group, the value of each replicate experimental wells minus background reactivity (i.e., three times the mean of the wells tested with pooled uninfected rabbit serum) was calculated and plotted. If residual value for the No-antigen control wells was present after subtraction, statistical significance was calculated with one-way ANOVA with the Bonferroni correction of multiple comparisons or t-test, with significance set at *p*<0.05. Except for figures showing cumulative absorbance, graphs represent the mean ± SEM for triplicate wells tested with pooled sera from the 3 rabbits in each group after background subtraction.

### TprC/D and D2 structure modeling

We used the ColabFold interface (58) to construct Multiple Sequence Alignments (MSA) for the TprC and TprD_2_ query sequences by searching UniRef30 (59), Mgnify (60) and ColabFold sequence databases with MMSeq2 (61). The MSA was used as input for structure prediction with AlphaFold2 (37) using the default settings (template=False, amber_relax=False, 3 recycles). Visualization was performed using PyMol software (https://pymol.org) (62).

## ACKNOWLEDGMENTS

Research reported in this publication was supported by National Institute of Allergy & Infectious Diseases of the National Institutes of Health under award number R01AI042143 grant (to SAL). Tpr models using AlphaFold2 were generated thanks to support from Open Philanthropy (to LG). This work was also partially supported also by the National Institute for Allergy and Infectious Diseases of the National Institutes of Health grant number U19AI144133 (Project 2. Project 2 leader: LG; PI: Anna Wald, University of Washington). The content is solely the responsibility of the authors and does not necessarily represent the official views of the Funders. The funders had no role in study design, data collection and interpretation, or the decision to submit the work for publication. The authors are grateful to Janelle Deane for aiding with some of the experimental procedures.

---

**Table S1.** TprC – IEDB B-cell epitope prediction results

| No. | Start | End | Peptide | Peptide(s) | Location | Length |
| --- | --- | --- | --- | --- | --- | --- |
| 1 | 41 | 59 | FQKNPRTGPGKHTHGFRIT | C2-C3 | ExL1 | 19 |
| 2 | 71 | 82 | KHTHTRRGARS | C5-C6 | Scaffolding | 12 |
| 3 | 95 | 113 | VELASSKSSTALSFTKPTA | C7-C9 | ExL2 | 19 |
| 4 | 132 | 180 | PSCVVNFAQLWKPFVTRAYSEKDTRYAPGFSGSGAKL<br>GYQAHNVGNSGV | C11-C16 | ExL3 | 49 |
| 5 | 192 | 208 | NGAWDSTDTHSKYGFG | C17-C18 | ExL4 | 17 |
| 6 | 215 | 220 | YGVDRQ | C19-C20 | Scaffolding | 6 |
| 7 | 233 | 251 | LDQNYVKGTEDSKNENKTA | C21-C23 | ExL5 | 19 |
| 8 | 277 | 295 | GNQHQSNAHAQTQERAILK | C26-C27 | ExL6 | 19 |
| 9 | 309 | 321 | QNLPNIMPPGIT | C29 | ExL6 | 13 |
| 10 | 343 | 352 | SAIQTVLAAG | C32-C33 | ExL6 | 10 |
| 11 | 359 | 377 | SQLVPNIEQGVDRVFRSSD | C34-C36 | ExL6 | 19 |
| 12 | 393 | 396 | PMNA | C37-C38 | ExL6 | 4 |
| 13 | 421 | 436 | TNIFGKRVFATTRAHY | C40 | ExL7 | 16 |
| 14 | 448 | 452 | KSGDP | C42-C43 | Scaffolding | 5 |
| 15 | 476 | 493 | FYRNNGGYELNRVPSGI | C45-C47 | ExL8 | 18 |
| 16 | 527 | 539 | NRFNIINAAGNLL | C50-C51 | ExL9 | 13 |
| 17 | 564 | 585 | WEQGVLSDPYMGIAESIWSER | C54-C56 | ExL10 | 22 |

**Table S2.** TprD_2_ – IEDB B-cell epitope prediction results

| No. | Start | End | Peptide | Peptide(s) | Location | Length |
| --- | --- | --- | --- | --- | --- | --- |
| 1 | 41 | 59 | FQKNPRTGPGKHTHGFRIT | C2-C3 | ExL1 | 19 |
| 2 | 71 | 82 | KHTHTRRGARS | C5-C6 | Scaffolding | 12 |
| 3 | 95 | 112 | VELASSKSSTALSFTKPT | C7-C9 | ExL2 | 18 |
| 4 | 132 | 180 | PSCVVNFAQLWKPFVTRAYSEKDTRYAPGFSGSGAK<br>LGYQAHNVGNSGV | C11-C16 | ExL3 | 49 |
| 5 | 192 | 208 | NGAWDSTDTHSKYGFG | C17-C18 | ExL4 | 17 |
| 6 | 215 | 220 | YGVDRQ | C19-C20 | Scaffolding | 6 |
| 7 | 233 | 255 | LDQNYVKGTEDSKNENKTALLWG | C21-C23 | ExL5 | 23 |
| 8 | 277 | 297 | GNQHQSNAQFYARMAPSQRVH | D26-D27 | ExL6 | 21 |
| 9 | 308 | 316 | LTSPQQDVV | D29 | ExL6 | 9 |
| 10 | 320 | 332 | VQELSKGSLLEKA | D30 | ExL6 | 13 |
| 11 | 339 | 362 | AQRTIVGLASSGGYLRHLNGKGLE | D32-D34 | ExL6 | 24 |
| 12 | 366 | 376 | RLIEQQKNPDA | D34-D35 | ExL6 | 11 |
| 15 | 419 | 434 | TNIFGERVFFKNQADH | D40 | ExL7 | 16 |
| 16 | 446 | 450 | KSGDP | C42-C43 | Scaffolding | 5 |
| 17 | 475 | 490 | YINNNGGAQYKGSNSDG | D45-D46 | ExL8 | 16 |
| 18 | 525 | 537 | NRFNHNQSGDALL | C50-C51 | ExL9 | 13 |
| 19 | 562 | 570 | WEQGVLAADA | C54-C55 | ExL10 | 9 |
| 20 | 578 | 583 | SIWSER | C56 | ExL10 | 6 |

**Table S3.** TprC – FBCPred B-cell epitope prediction results

| Position | Epitope | Peptide(s) | Location | Score |
| --- | --- | --- | --- | --- |
| 42 | QKNPRTGPGKHTHG | C2-C3 | ExL1 | 0.999 |
| 70 | SKHTHTRRGEARSG | C5-C6 | Scaffolding | 0.895 |
| 105 | ALSFTKPTASFQAT | C8-C9 | ExL2 | 0.962 |
| 130 | TSPSCVVNFAQLWK | C11-C12 | ExL3 | 0.985 |
| 156 | RYAPGFSGSGAKLG | C14-C15 | ExL3 | 0.998 |
| 171 | QAHNVGNSGVDVDI | C15-C16 | Scaffolding | 0.992 |
| 191 | SNGAWDSTDTHSK | C17-C18 | ExL4 | 0.995 |
| 241 | TEDSKNENKTALLW | C22-C23 | ExL5 | 0.923 |
| 259 | RLTLEPGAGFRFSF | C24-C25 | Scaffolding | 1 |
| 308 | VQNLPNIMMPPGIT | C29 | ExL6 | 0.809 |
| 362 | VPNIEQGV RDVFRS | C34-C35 | ExL6 | 0.935 |
| 441 | GFLKLETSGDPYT | C42-C43 | Scaffolding | 0.997 |
| 464 | VEARVYIPLTYV FY | C45 | ExL8 | 0.825 |
| 524 | GTTNRFNIINAAGN | C51-C52 | ExL9 | 0.996 |
| 561 | SAQWEQGVLS DVPY | C54-C55 | ExL10 | 0.919 |

**Table S3.** TprD_2_ – FBCPred B-cell epitope prediction results

| Position | Epitope | Peptide(s) | Location | Score |
| --- | --- | --- | --- | --- |
| 42 | QKNPRTGPGKHTHG | C2-C3 | ExL1 | 0.999 |
| 70 | SKHTHTRRGEARSG | C5-C6 | Scaffolding | 0.895 |
| 105 | ALSFTKPTASFQAT | C8-C9 | ExL2 | 0.962 |
| 130 | TSPSCVVNFAQLWK | C11-C12 | ExL3 | 0.985 |
| 156 | RYAPGFSGSGAKLG | C14-C15 | ExL3 | 0.998 |
| 171 | QAHNVGNSGVDVDI | C15-C16 | Scaffolding | 0.992 |
| 191 | SNGAWDSTDTHSK | C17-C18 | ExL4 | 0.995 |
| 241 | TEDSKNENKTALLW | C22-C23 | ExL5 | 0.923 |
| 259 | RLTLEPGAGFRFSF | C24-C25 | Scaffolding | 1 |
| 308 | LTSPQQDVVSFFVQ | D29 | ExL6 | 0.8 |
| 365 | MRLIEQQKNPDARM | D34-D35 | ExL6 | 0.829 |
| 439 | GFLKLETSGDPYT | D42 | Scaffolding | 0.997 |
| 462 | VEARVYIPLTYIFY | D45-D46 | ExL8 | 0.975 |
| 483 | YKGSNSDGVINTPI | D47 | ExL8 | 0.996 |
| 523 | ATNRFNHNQSGDAL | D51-D52 | ExL9 | 0.956 |
| 558 | LSAQWEQGV LADAP | C54-C55 | ExL10 | 0.895 |

**Fig. S1.**
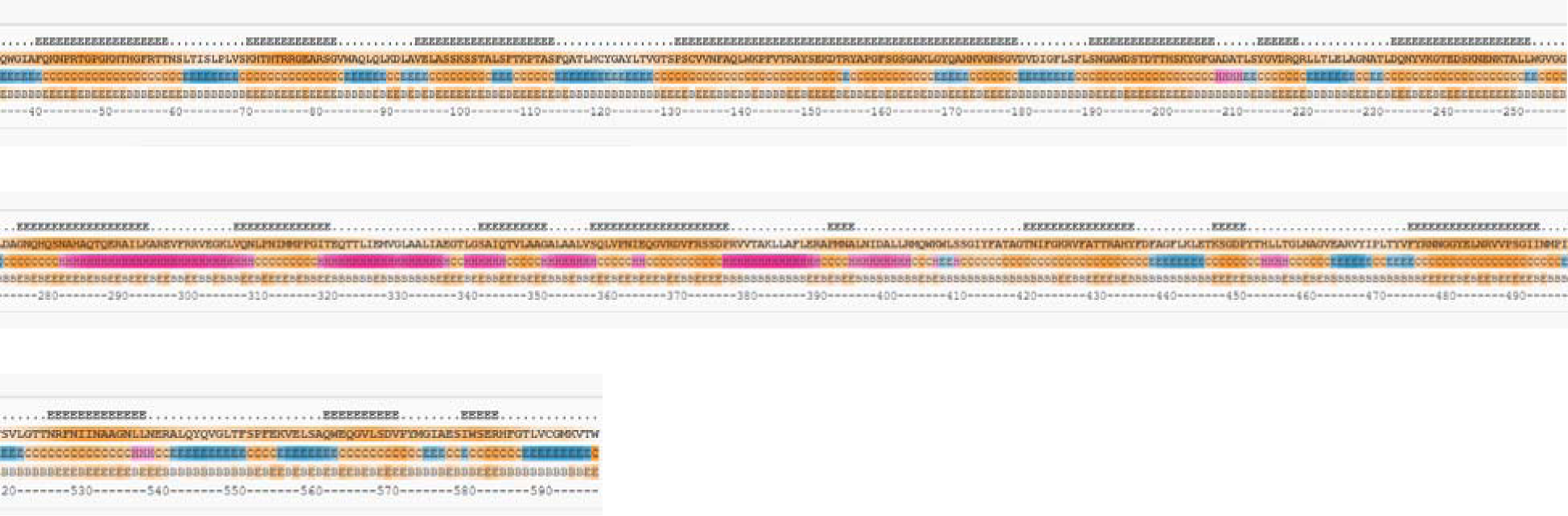
TprC- BepiPred B-cell epitope prediction results.

**Fig. S2.**
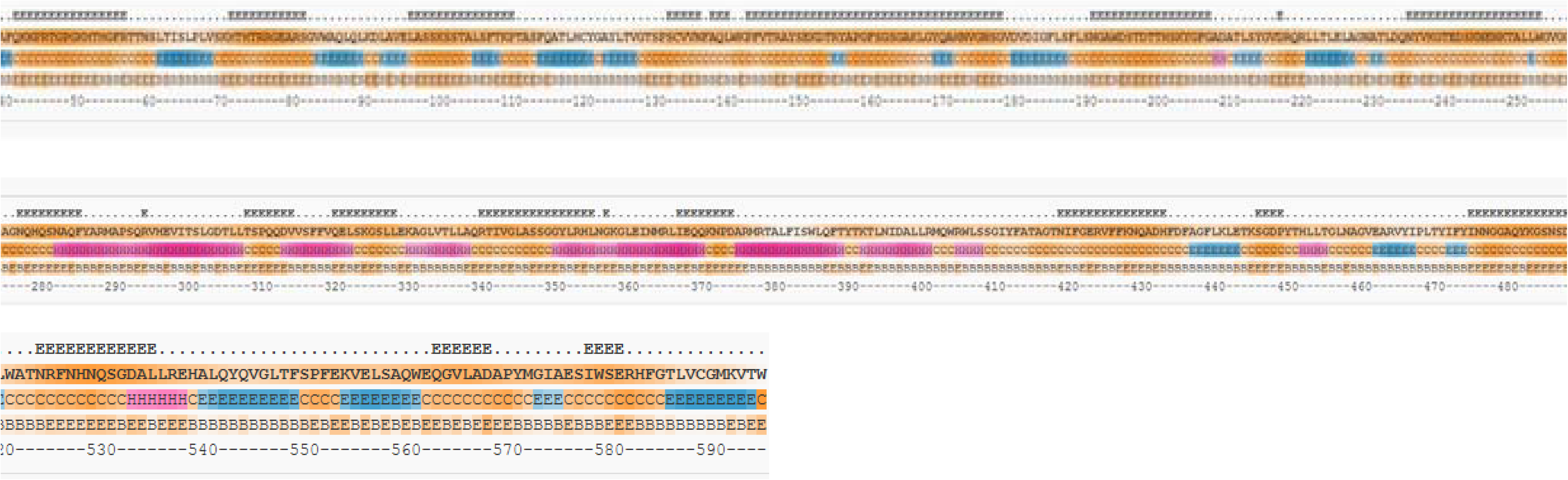
TprD_2_- BepiPred B-cell epitope prediction results.

**Fig.S1/2 Legend.**
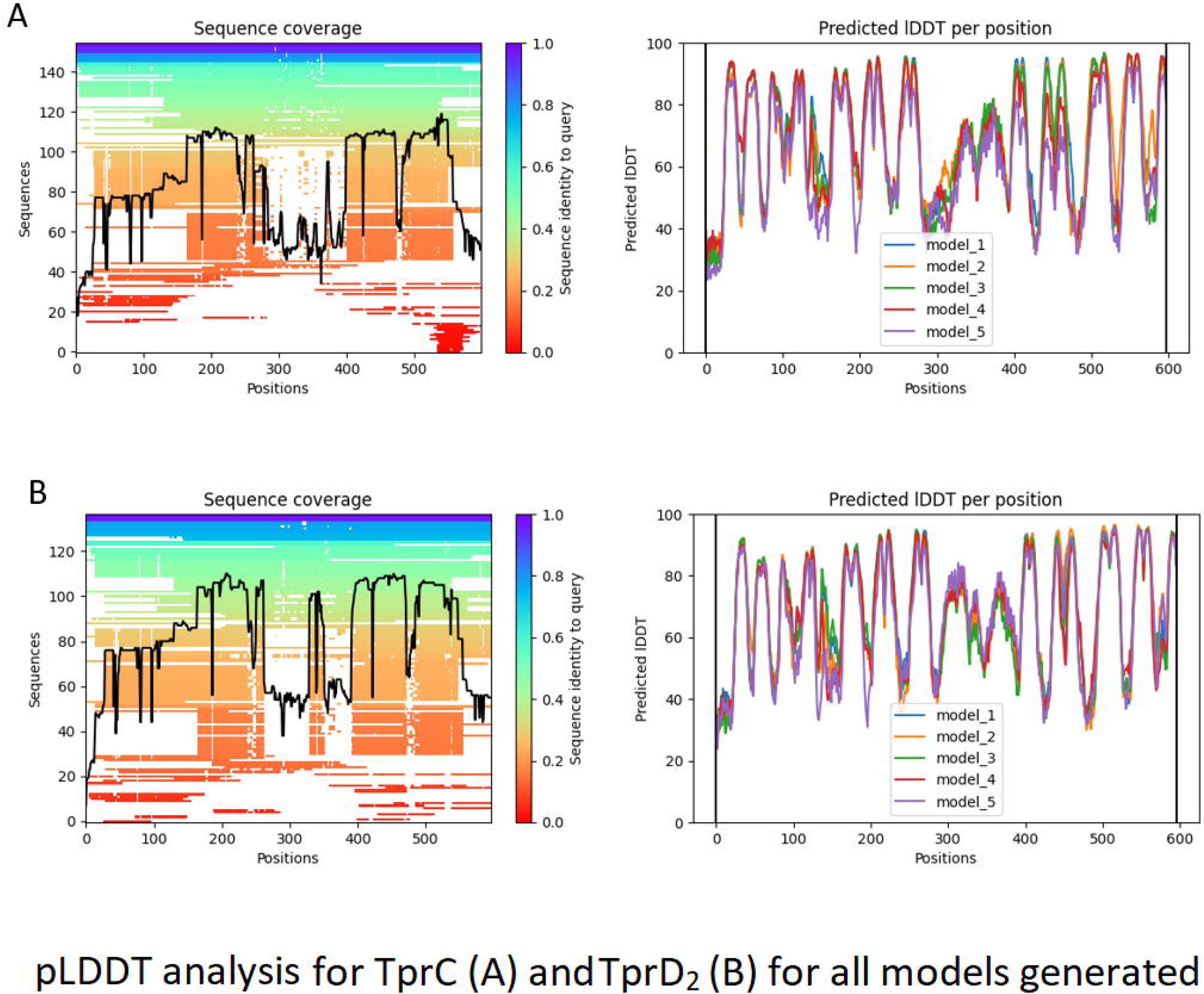
The output format shows the BepiPred-2.0 predictions and epitope classification for each sequence. The BepiPred-2.0 predictions are used to set the background color of the protein sequences. All predictions greater than a user-defined threshold (by default 0.5) are marked as ‘E’ in the ‘Epitopes’ line above the protein sequence itself.

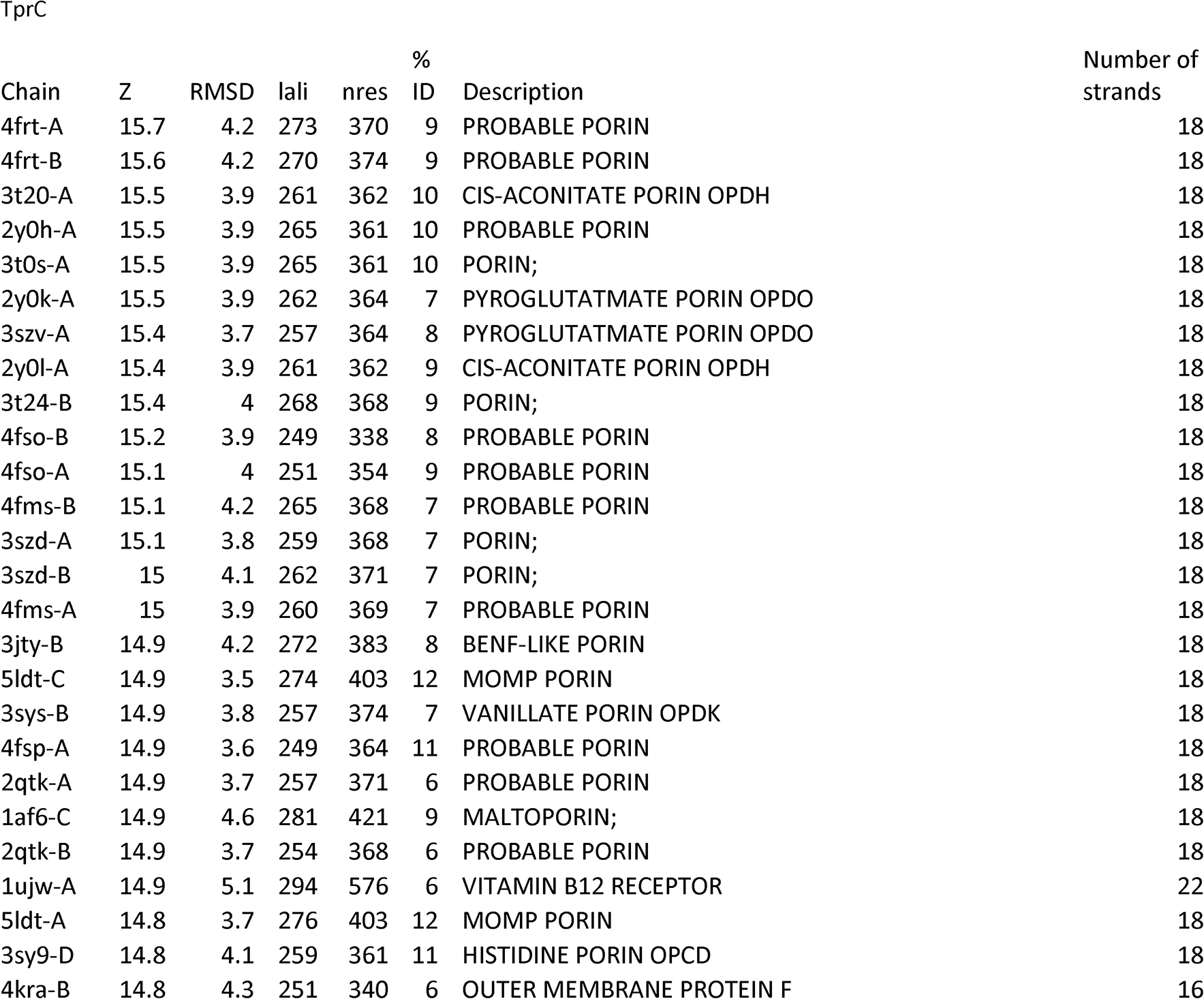
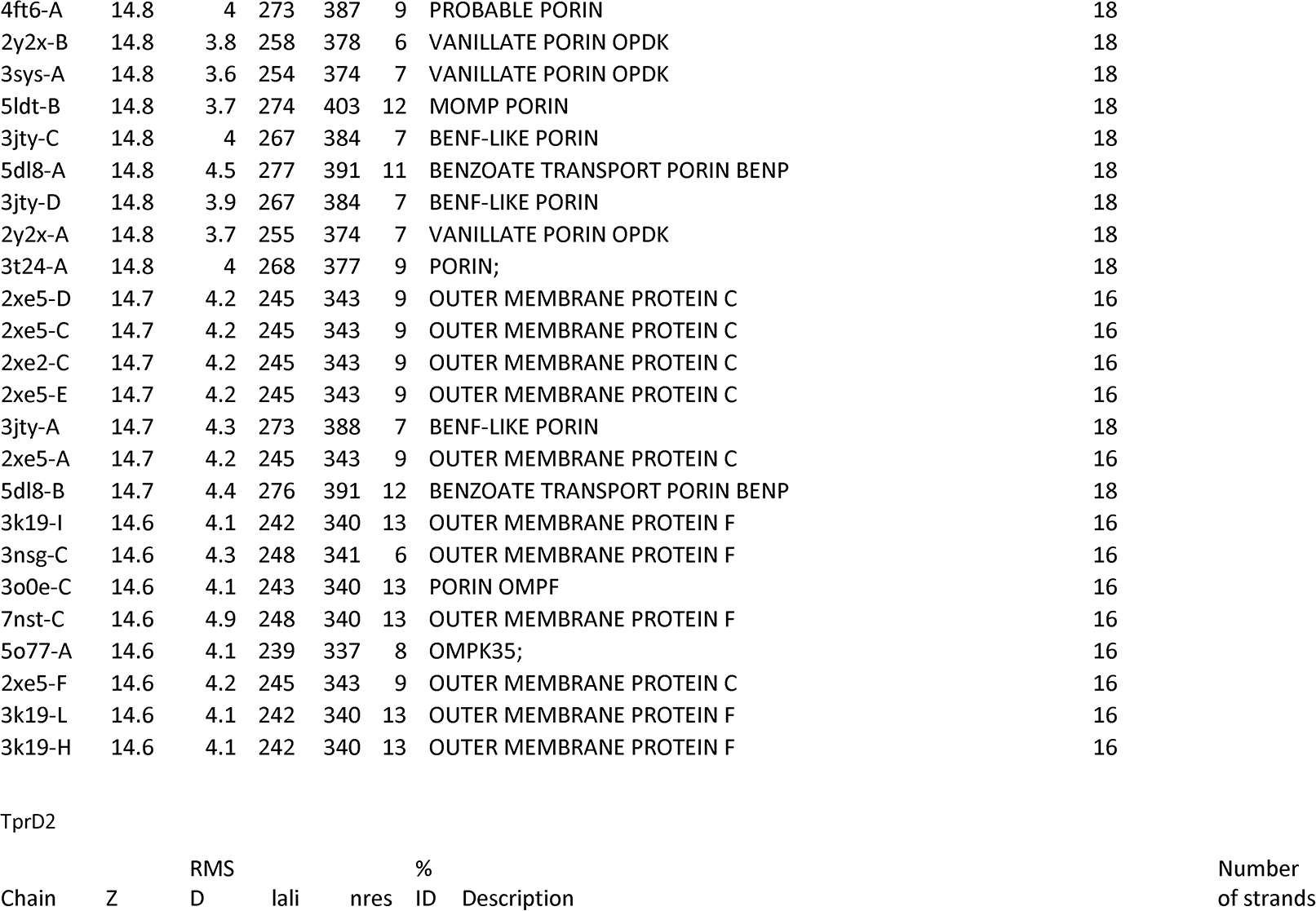
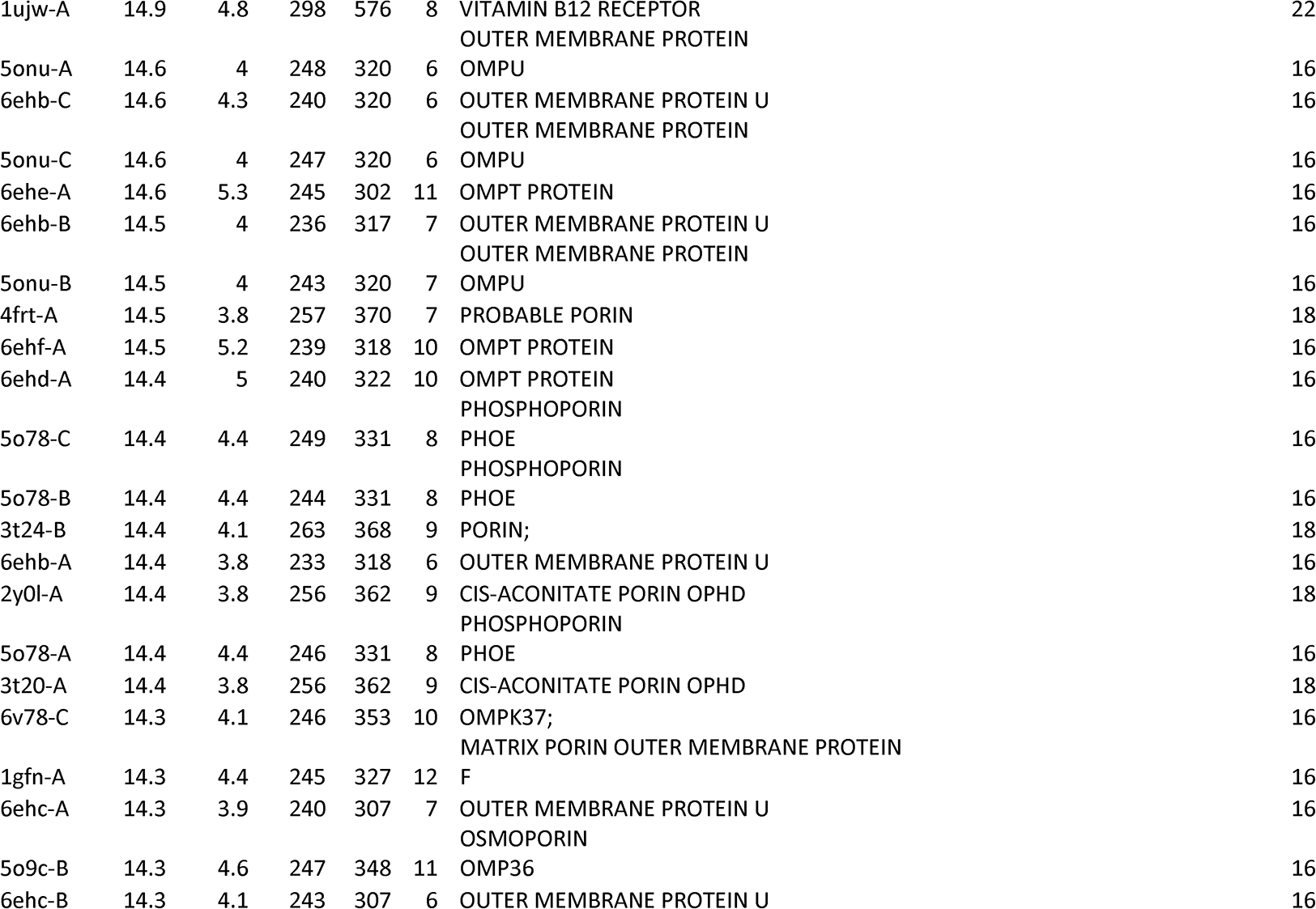
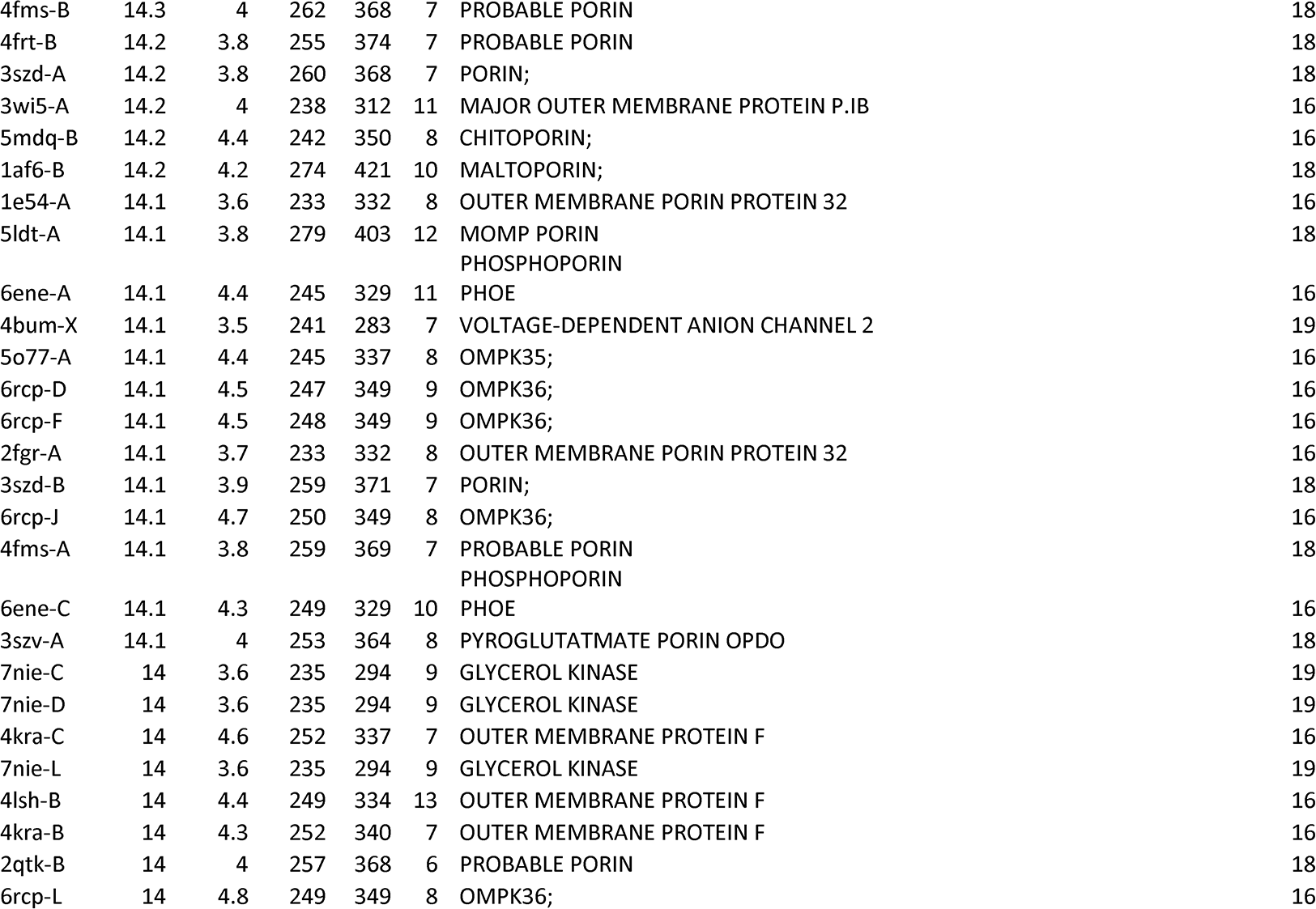
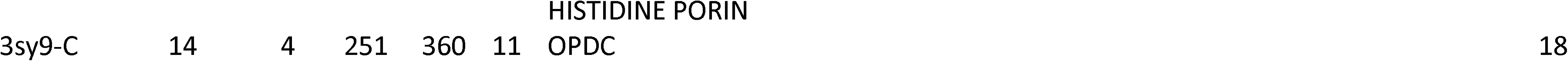

